# Multilayer and Biomimetic Polycaprolactone/Uterine ECM Scaffold for Uterine Tissue Engineering and Anti-Adhesion Barrier Applications via 3D-Printing Near-Field Melt Electrowriting

**DOI:** 10.64898/2026.09.13.751266

**Authors:** Abbas Fazel Anvari-Yazdi, Kobra Tahermanesh, Maryam Ejlali, Daniel J. MacPhee, Ildiko Badea, Xiongbiao Chen

## Abstract

**Background:** Post-operative uterine adhesions remain a major clinical challenge, often leading to infertility and severe pelvic pain. Current anti-adhesion barriers lack the mechanical strength, tissue specificity, or bioactivity required for optimal regeneration of uterine tissue. To address these limitations, we developed a biomimetic, multilayer scaffold that replicates the structural and functional complexity of the myometrial and serosal layers of the uterus by combining a tunable polycaprolactone (PCL) mesh with a bioactive hydrogel derived from decellularized uterine extracellular matrix (dUECM) reinforced with sodium alginate (Alg).

**Materials and Methods:** Composite scaffolds were fabricated using near-field melt electrowriting (MEW) to create precision-engineered PCL meshes with various infill angles (90°, 60°, 45°, and 30°), followed by impregnation with an alginate-dUECM hydrogel. A comprehensive physicochemical characterization was conducted using Fourier Transform Infrared Spectroscopy (FTIR), Thermogravimetric Analysis (TGA), and contact angle measurements. Mechanical properties were evaluated in both dry and wet states via uniaxial tensile testing. Swelling and degradation were assessed over 10 days. hTERT-HM cells were used for in vitro evaluation through Live/Dead staining, MTT, SEM, and immunocytochemistry for α-SMA and DAPI.

**Results:** The successful incorporation of the hydrogel into the PCL mesh was confirmed by the results obtained from FTIR and TGA analyses; these findings were further supported by contact angle studies, which showed a marked increase in hydrophilicity, resulting in a reduction of the static contact angle from 113 ± 4° in the pure PCL to 60 ± 12° in the hydrogel-infiltrated samples in the 30° group. Mechanical testing showed that the wet composite scaffolds maintained higher stiffness than hydrated native uterine tissue. Cell culture assays confirmed excellent cytocompatibility and proliferation, especially in 30° infill architecture. Immunofluorescence revealed strong α-SMA expression and cytoskeletal organization, indicating phenotypic maturation of uterine smooth muscle cells for all groups.

**Conclusion:** Among all groups, the 30° MEW mesh impregnated with Alg-dUECM hydrogel demonstrated the most balanced combination of mechanical resilience, degradation profile, wettability, and cellular compatibility. This structure most closely recapitulates the biological characteristics of uterine outer myometrium tissue, positioning it as a highly promising scaffold for regeneration and biofunctional anti-adhesion barrier applications.

## 1. Introduction

Post-operative adhesions—fibrous bands that abnormally link organs after surgery—develop in up to 90 % of open abdominal or pelvic procedures and remain a leading cause of chronic pain, bowel obstruction, and infertility, especially after gynaecologic surgery [1, 2]. The most effective preventive strategy is a temporary physical barrier that separates injured tissue planes during the first 3–7 days of wound healing, when fibrin deposition and fibroblast proliferation drive adhesion formation [3]. Ideally, such a barrier would also promote regenerative repair rather than merely block contact. Achieving this dual function demands materials that are simultaneously anti-adhesive, bioactive, and mechanically supportive—attributes seldom met by a single component. A synergistic solution is to pair a bioactive material, such as a uterine-specific hydrogel derived from decellularized extracellular matrix, with a structurally robust, slowly resorbing synthetic polymer.

Among the available biomaterials, decellularized uterine extracellular matrix (dUECM) has emerged as a promising bioactive option due to its native biochemical composition and inherent signaling capacity, which support cellular functions and tissue integration [4, 5]. Unlike inert physical barriers, dUECM-based hydrogels or films can promote regenerative healing while still providing temporary separation of tissue planes [6]. When combined with US FDA-approved synthetic polymers such as poly(ε-caprolactone), a versatile, biodegradable polyester known for its mechanical strength and slow degradation profile, hybrid scaffolds can be engineered to achieve a balance between bioactivity and structural integrity [7–9]. The fabrication of dUECM-based hydrogels and their integration into multilayered designs allows for enhanced tissue mimicry, while PCL offers the precision required for architectural reinforcement. Notably, PCL can be processed using advanced manufacturing methods such as 3D printing, electrospinning, and melt electrowriting (MEW) [10]. MEW offers unparalleled control over micro-scale fiber alignment and scaffold geometry.

MEW is a precise form of melt electrospinning that enables the fabrication of micro fiber scaffolds with controlled architectures. The concept dates back to early melt electrospinning studies in 1981, but modern MEW was established around 2011 as a direct writing technique for depositing fibers in defined patterns [11, 12]. In MEW, a polymer (commonly thermoplastic polyesters like PCL) is melted and extruded through a fine nozzle while a high voltage is applied between the nozzle and a grounded collector [13]. MEW yields open, highly porous meshes with user-defined fiber spacing and orientation, enabling precise control of scaffold anisotropy while avoiding solvent residues and random lay-down inherent to solution electrospinning [14–16].

This manufacturing precision is vital because the uterine wall is strongly layer-dependent. The outer myometrium contains longitudinally oriented smooth-muscle bundles and collagen fibres that generate contractile force, whereas the serosa consists of a mesothelial cell sheet supported by loosely arranged, surface-parallel ECM that minimises friction with adjacent organs [17–19]. Replicating this anisotropic–isotropic gradient is essential for restoring uterine biomechanics and for preventing the very adhesions the barrier must block. Achieving seamless multilayer assembly (myometrium vs. serosa), ensuring vascularization, and accommodating uterine expansion and movements remain critical engineering challenges that must be overcome for any scaffold intended to function safely and effectively in vivo [20].

Therefore, this research aims to develop a hybrid and functional scaffold by integrating the architectural advantages of MEW-fabricated PCL microstructures with the biochemical and biological cues provided by dUECM-derived hydrogels. Through a series of physicochemical characterizations and in vitro biological assays, this study seeks to optimize scaffold composition, mechanical performance, and cellular responses—paving the way for a scalable, bioinspired platform tailored for uterine tissue regeneration and post-operative anti-adhesion applications.

## 2. Materials and Methods

### 2.1. Decellularization of Porcine Uterine Tissue

Uterine tissue was decellularized according to our previously published one-step Triton X-100/SDS method with no modifications [4, 21]. Briefly, fresh porcine uterine tissue was collected from a 1-year-old pig (∼200 kg) and transported to the lab in sterile phosphate buffer saline (PBS; Sigma-Aldrich, Cat.# P4417) within 30 min. Uteri were frozen at −80 °C, then sectioned into ∼2 mm discs and thawed overnight (4□°C). Discs (30 g/500 mL) were pre-rinsed for 24 h in Dulbecco’s Phosphate Buffered Saline (DPBS; Gibco, Cat.# 21600) containing 1 % antibiotic/antimycotic (Gibco, Cat.# 15240062) on an orbital shaker at 200 rpm, 4□°C. Decellularization was performed for 48 h in a single bath of 1 % SDS (Sigma-Aldrich, Cat.# L3771) + 1 % Triton™ X-100 (Fisher Bioreagents, Cat.# BP151), with fresh reagent every 8 h. Residual detergents were removed by washing in cold Milli-Q water for 72 h (changes every 8 h) until no foaming was observed by shaking. Once the tissue was decellularized, it was freeze-dried to remove residual moisture, thereby enhancing its stability and shelf life.

### 2.2. Near-Field Melt Electrowriting (NF-MEW) Scaffold Fabrication

Poly(ε-caprolactone) (PCL) (Mn 45,000, Sigma-Aldrich, Cat.# 704105) was used for melt electrowriting (MEW) scaffold fabrication. The PCL was processed without additional purification. All scaffolds were fabricated using a GeSIM BioScaffolder 3.1 Melt Electrowriting system equipped with a heated stainless steel nozzle (250 µm inner diameter). The polymer melt was extruded using nitrogen (N ) pressure, which provided a controlled and stable flow of molten PCL. The melted polymer was directed onto a grounded spinneret positioned over a glass tray, with a high-voltage (HV) power supply connected to the collector tray to facilitate electrostatic fiber deposition. The PCL was melted at 100°C and maintained at this temperature throughout the printing process. Printing was performed under ambient conditions (∼22°C, standard laboratory humidity).

The fabrication of architecturally defined scaffolds is predicated on precise control over the fundamental physics of the MEW process, which is depicted schematically in Fig. 1A. The method utilizes a programmable 3-axis gantry equipped with a high-temperature spinneret, which N_2_ pressurizes to extrude molten PCL. A high-voltage electric field (5 kV) established between the charged spinneret and a collector deforms the polymer melt into a Taylor cone. This initiates a stable, charged jet that is drawn toward the collector, solidifying into a fine strand/fiber.

**Figure 1:**
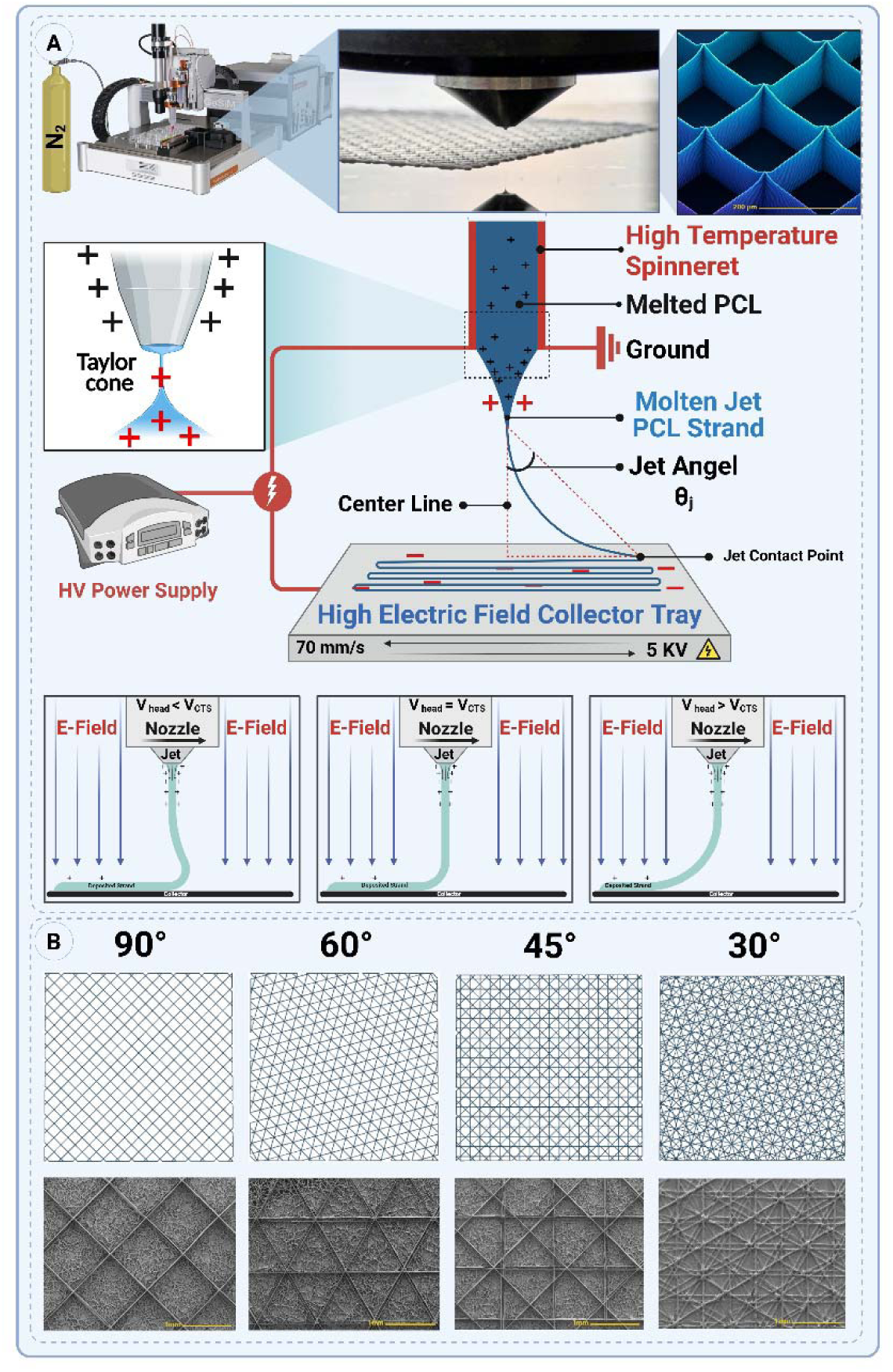
Near-field melt-electrowriting (NF-MEW) set-up and architectural tuning of PCL meshes. (A) A heated spinneret (100 °C) extrudes molten PCL through a Taylor cone, while a +5 kV potential is applied across a 2 mm gap to the collector. The print-head travels at 70 mm s^−^¹, imposing a controllable jet angle θ^−^ relative to the centre-line. Insets illustrate how the jet path evolves when the head speed is below, equal to, or above the critical translation speed (V_CTS_). Photographs (top right) show a multilayer mesh forming in real time and a SEM close-up of the resulting 200 µm pores. (B) Designed mesh geometry as a function of raster angle. Plan-view CAD patterns (top row) and corresponding SEM micrographs (bottom row) for bilayer grids printed with inter-strand angles of 90°, 60°, 45° and 30°. Decreasing the raster angle increases filament density and transforms square pores into equilateral, rhombic or star-like motifs while retaining straight, defect-free strands at the optimized process window (70 mm s^−^¹, 3 kPa, 5 KV, IF 2mm, Temp. 100 °C).

High fidelity can only be achieved with a complete understanding of the complex relationship between the mechanical motion of the head and the electrostatic forces that affect the jet. To optimize the MEW printing of PCL strands, we conducted printability experiments using the parameters listed in Table 1.

**Table 1:**
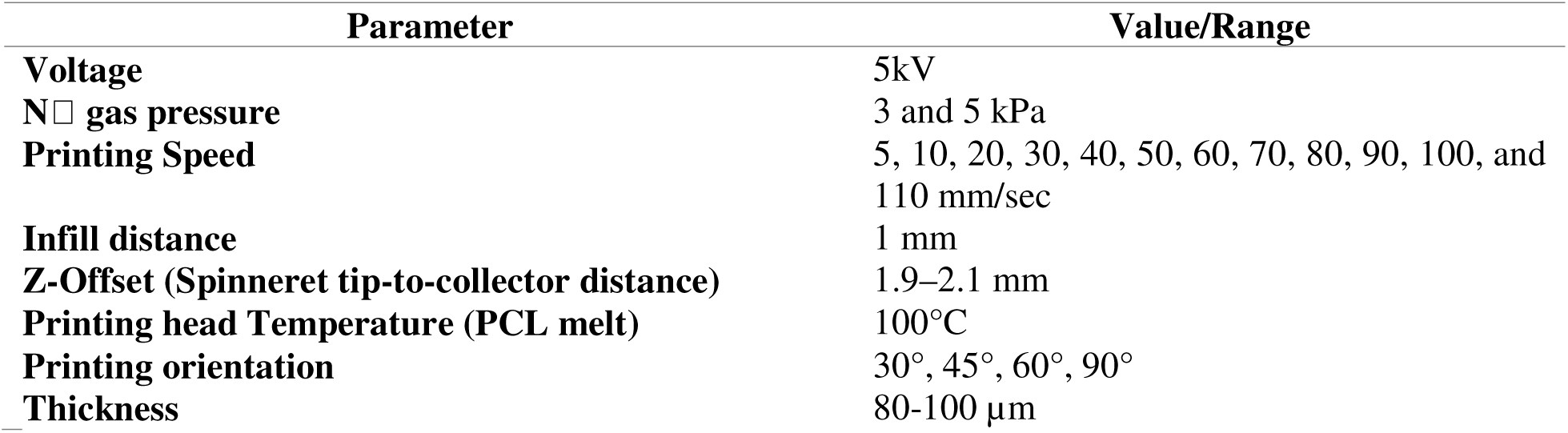
MEW printing parameters.

Following fabrication, the scaffolds were examined under an optical and scanning electron microscope (SEM) to assess fiber alignment and uniformity.

### 2.3. Hydrogel ink preparation

Alginate and dUECM were the primary components used to prepare hydrogel ink. The dried dUECM was subsequently ground into a fine powder to facilitate enzymatic digestion. To achieve this fine particulate form, the freeze-dried tissue was pulverized using dry ice and a grinder to ensure uniformity and minimize aggregation. This step is critical for enhancing the reactivity of the powder in subsequent digestion.

The required quantity of dUECM powder was weighed and then subjected to enzymatic digestion. To ensure optimal digestion, the Voytik-Harbin method was used [22]. An aliquot of 100 mg of dUECM powder was suspended in a 0.5 M acetic acid solution (Cat.# 351271, Fisherbrand) containing 10 mg of Pepsin (Cat.# P7125, Sigma-Aldrich) to catalyze the digestion process. The suspension was maintained at a temperature of 4°C under constant stirring (50 rpm) for 24 hours to ensure complete digestion. The cold environment helped preserve the structural integrity of the ECM components during the enzymatic reaction.

Following digestion, the resulting dUECM solution was neutralized with 10M sodium hydroxide (NaOH, Cat. # BP359, Fisher Scientific) [23]. This neutralization step was performed in a refrigerated environment at 4°C within a walk-in cooling chamber to prevent premature self-crosslinking of collagen molecules, which could compromise the hydrogel’s handling and mixing with sodium alginate. The slow and controlled neutralization allowed for the formation of a homogeneous and stable hydrogel suitable for further steps.

To ensure the removal of residual components from the neutralization reaction (such as sodium ions and unreacted NaOH) as well as pepsin, the neutralized dUECM solution was subjected to dialysis. A 12–14 kDa molecular weight cut-off (MWCO) membrane was used to retain larger ECM components such as collagen and glycosaminoglycans while allowing smaller molecules like sodium ions, hydroxide ions, and pepsin to diffuse out.

The solution was dialyzed against a neutral PBS buffer. The dialysis process was carried out at 4°C to prevent degradation of ECM components, with buffer changes performed multiple times over a 48-hour period. This protocol ensured thorough removal of small molecules and enzymes while preserving the integrity and bioactivity of the ECM components.

#### 2.3.1. Alg-dUECM hydrogel preparation and scaffold loading

Following our optimization studies, a 3 % (w/v) alginate and 1 % (w/v) dUECM hydrogel formulation was selected for mesh impregnation. Sodium alginate was dissolved in molecular biology-grade water (Hypure; Cytiva, Cat.# 82007-334) and subsequently mixed with dUECM under sterile conditions in a biosafety cabinet. All preparation steps were conducted on ice to preserve hydrogel integrity and prevent premature gelation. The prepared hydrogel was then centrifuged at 2,000 rpm for 15 minutes at 4□°C using a pre-cooled centrifuge to remove entrapped air bubbles and ensure homogeneous consistency prior to scaffold application.

#### 2.3.2. Impregnation and cross-linking

MEW-PCL meshes (on glass coverslips) were placed in 6-well plates; 250 µL of cold Alg-dUECM hydrogel was pipetted onto each construct and the plate centrifuged (4□°C, 10 min) to force hydrogel penetration. Constructs were ionically cross-linked in 50 mM CaCl (Cat.# 223506, Sigma-Aldrich), rinsed thoroughly with cold, fully supplemented DMEM/F12 medium (Cat.# 11320033, Gibco), and then incubated at 37 °C and 5% CO for 6 h to allow additional ECM self-assembly. Fully gelled samples were frozen at −20 °C overnight and freeze-dried.

#### 2.3.3. Secondary PCL reinforcement

Freeze-dried hydrogels were returned to the MEW collector and sealed with two randomly oriented PCL fibre layers to stabilise the aerogel core.

#### 2.3.4. hTERT-HM cell embedding

Cells were detached with 0.25 % Trypsin-EDTA (GE HyClone, Cat.# SH30042.02), neutralised in DMEM/F-12 (Gibco, Cat.# 11320033) containing 10 % FBS (Cytiva, Cat.# CA76236-336), pelleted (1000 rpm, 5 min), and washed twice in fully supplemented DMEM/F12 media. A suspension of 2 × 10□ cells mL^−^¹ was prepared on ice-cold 3 % Alg + 1 % dUECM hydrogel; ∼5 × 10□ cells (250 µL) were gently poured onto each scaffold before culture in complete medium.

### 2.4. Scanning electron microscopy (SEM)

The microstructure of prepared MEW meshes were investigated using field-emission scanning electron microscopy (FE-SEM; Hitachi SU8010). For samples seeded with hTERT-HM cells, they were fixed using 2.5% glutaraldehyde (GA; Polysciences Inc., Cat.# 00376) at 4°C overnight, then gradually dehydrated by an increasing concentrations of ethanol (10%, 30%, 60%, 90%, and 100%) [24, 25]. After dehydration, the samples were immersed into 1:2 and 2:1 hexamethyldisilizane (HMDS; Thermo Scientific, Cat.# A15139. AP): absolute ethanol, respectively, for 20 min and then 100% HMDS solution overnight. Samples were air-dried in a fume hood. Eventually, specimens were mounted on aluminum stubs with double-sided carbon tape and coated with 10 nm of gold (Quorum Q150TES Sputter Coater) as the conductive layer. For imaging, the voltage was fixed at an accelerating voltage of 3 KV to capture the secondary electron mode images.

### 2.5. FT-IR spectroscopy

IR spectra of the specimens were acquired using Attenuated Total Reflectance-Fourier Transform Infrared spectroscopy (ATR-FTIR; Spectrum 3 Tri-Range MIR/NIR/FIR Spectrometer, PerkinElmer). The specimens were placed directly on the ATR crystal *(n = 3).* FT-IR spectra were acquired in 4,000–600 cm^−1^ range at a resolution of 4 cm^−1^.

### 2.6. Thermogravimetric analysis (TGA)

To assess the PCL/Alg/dUECM decomposition behaviour, thermogravimetric analysis (TGA) was performed using a Perkin-Elmer TGA 8000 analyzer. Approximately 10 mg of the prepared samples underwent heating from 50°C to 900°C at a rate of 10°C/min under a constant nitrogen gas flow of 30 cm^3^/min. By differentiating the TGA values, the differential form of TGA (DTA) was derived, facilitating the identification of the maximum disintegration temperature at each stage of thermal degradation.

### 2.7. Swelling and degradation

To evaluate the swelling and degradation characteristics of 3D-printed alginate-dUECM scaffolds, the samples were incubated in complete DMEM/F12 culture media at 37°C under 5% CO_2_. The scaffolds were fabricated under sterile conditions to prevent contamination that might influence swelling and mass loss measurements.

The initial dry weight of each sample was recorded as W . At predetermined time points (0, 1, 2, 6, 12, and 18 hours), the scaffolds were gently blotted with sterile Kimwipes™ to remove surface moisture, and the corresponding wet weight (W ) was measured. The swelling ratio was then calculated using the following [26]:

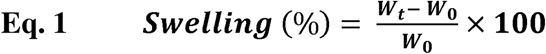

Measurements were conducted in quintuplicate for each time point to ensure statistical accuracy.

Following the swelling assessments, the scaffolds were freeze-dried to determine their dry weight at specific intervals (*W_l_*). The remaining mass percentage was calculated using the formula [26]:

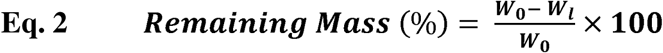

Where *W_0_* is the initial dry weight of the scaffold, and *W_l_* is the lyophilized weight at each time point. This process was performed for each interval (0, 2, 4, 7, and 10 days) using separate sets of scaffolds to maintain accuracy. All experiments were conducted in sterile conditions, and results were analyzed to determine the swelling and degradation behaviors of the scaffolds over time.

### 2.8. Cell viability for developed hydrogel ink (MTT and Live/Dead Assays)

Cell viability of the MEW scaffolds were assessed using MTT (3-[4,5-dimethylthiazol-2-yl]-2,5-diphenyltetrazolium bromide, Cat. # M6494, Invitrogen) and Live/Dead assays. To evaluate the metabolic activity of hTERT-HM cells seeded on the scaffolds, MTT assays were performed on days 1, 5, and 7. MTT solution (0.5 mg/mL) was added to each well and incubated for 4 hours at 37°C, allowing metabolically active cells to form formazan crystals. These crystals were dissolved in DMSO (Dimethyl sulfoxide, Cat. # BP231-1, Fisher bioreagents) and absorbance was measured at 550 nm using a microplate reader (BioTek Synergy HT Multi-Detection Microplate Reader) to assess cell proliferation and viability over time.

The live/dead assay was performed to evaluate the viability, attachment, and distribution of hTERT-HM cells on the scaffolds. Calcein AM (Cat. # AS-89201, AnaSpec, Fremont) and propidium iodide (PI, Cat. # AS-83215, AnaSpec, Fremont) were used to stain live and dead cells, respectively. Imaging was performed using a Lionheart Biotek fluorescence microscope in stacked imaging mode to minimize blurring caused by surface irregularities. The images were deconvoluted using the microscope’s software to enhance clarity and analyze cell viability and morphology.

### 2.9. Tensile testing of decellularized tissues

Tensile properties of both dry and hydrated MEW scaffolds were evaluated, with native porcine myometrium serving as the biological control. All samples were prepared in a rectangular shape with consistent dimensions. Testing was performed using an Electroforce BioDynamic 5110 mechanical tester (TA Instruments). The crosshead speed was set to 0.01 mm/sec, which is appropriate for soft tissue characterization. Stress–strain curves were generated from the force–displacement data, and key mechanical parameters including ultimate tensile strength (UTS), Young’s modulus, and elongation at rupture (%) were calculated.

### 2.10. Wettability Measurements

Wettability of the MEW scaffolds was assessed using a static contact angle measurement setup custom-designed in the laboratory. A 4 μL droplet of pre-warmed complete DMEM/F-12 medium was gently deposited onto the scaffold surface using a micropipette. Contact angle measurements were recorded using a high-speed camera positioned perpendicularly to the scaffold surface. The initial static contact angle was measured immediately upon droplet placement. Furthermore, we measured the absorption time, which is defined as the time between a droplet’s deposition and its complete absorption into the scaffold, in order to assess the scaffold’s liquid uptake characteristics. To ensure accuracy and reproducibility, all measurements were meticulously conducted at a consistent room temperature, under standardized lighting conditions, and within a carefully controlled environmental setting.

### 2.11. Immunofluorescence (IHC)

MEW scaffolds were fixed in 4% paraformaldehyde (PFA) in PBS and stored at 4□°C. For immunocytochemistry, scaffolds were washed in PBS for 5 minutes, then permeabilized using PBS with 0.1% Triton X-100 (PBT) for 15 minutes at room temperature (RT) on a shaker. After an additional PBS wash, blocking was performed for 30 minutes at RT in a solution of 5% goat serum (Vector Laboratories, VECTS1000), 1% horse serum, and 1% FBS in PBS.

Scaffolds were then incubated overnight at 4□°C with primary antibodies diluted in blocking buffer (e.g., mouse IgG at 10□μg/mL), placed face-down on 200□μL droplets of antibody solution on Parafilm inside a humidified chamber. Following the initial procedures, the samples were subjected to three washes using PBT, after which they underwent a one-hour incubation period at room temperature in the dark with secondary antibodies (anti-mouse RRX, diluted 1:150 in blocking buffer). This was followed by three washes in PBT and two final PBS washes, each for 5 minutes.

Finally, scaffolds were kept in PBS at 4□°C, protected from light, until confocal imaging. Images were acquired using a Nikon Eclipse Ti2-E fluorescence microscope.

### 2.12. Statistical Analysis

The normality of data distribution was assessed using the Shapiro-Wilk test, and appropriate statistical tests were selected based on the results. Data are presented as mean ± SD. Statistical significance was determined using one-way or two-way ANOVA with Tukey’s post hoc test. The significance levels are indicated as follows: **** for p < 0.0001, *** for p < 0.001, ** for p < 0.01, * for p < 0.05, and NS for non-significant differences.

## 3. Results

### 3.1. Thermal profile of the PCL and *Melt Electrowriting Printability*

The primary objective of this study was to establish a robust and high-fidelity NF-MEW process for fabricating PCL scaffolds with pre-designed micro-architectures. This was achieved through a two-stage optimization process: first, a broad-range screening of printing speed to identify a promising processing window, followed by a targeted refinement of pressure and speed to pinpoint the optimal set of parameters.

Initially, scaffolds were printed at a constant pressure of 5 kPa while the printing speed was systematically varied from 5 to 110 mm/s. The material’s thermal properties were confirmed by DSC, showing a sharp crystallization temperature of 89.14□°C, ensuring rapid fiber solidification upon deposition (Fig. 2B). Qualitative analysis revealed three distinct printing regimes (Fig. 2A). At low speeds (5-20 mm/s), the process was dominated by jet instability, resulting in severe coiling and looping of the polymer fiber. This completely obscured the intended grid-like architecture, producing a non-uniform fibrous mat instead of an ordered strand. As speed increased (30-70 mm/s), the coiling instability was suppressed and replaced by a wavy or meandering fiber deposition. The strands’ grid structure became progressively more apparent, but the fibers lacked the straightness required for high fidelity. A stable, high-fidelity regime was achieved at speeds of 80 mm/s and above. The drawing forces exerted by the fast-moving head effectively linearized the polymer jet, resulting in the deposition of perfectly straight and continuous fibers. The scaffolds fabricated in this regime exhibit exceptional geometric precision, accurately replicating the programmed design.

**Figure 2:**
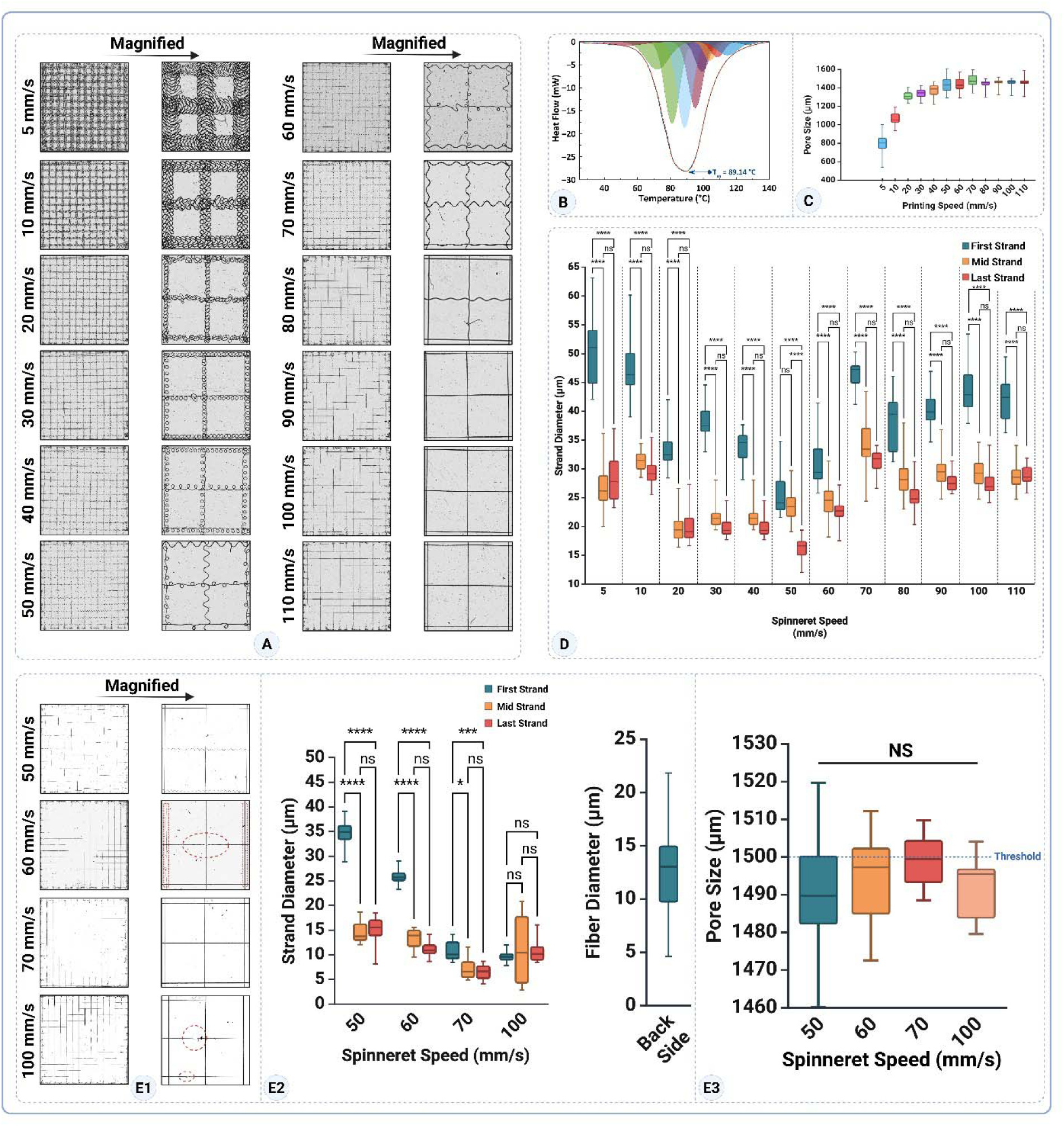
Printability window and optimization of PCL melt-electrowriting. (A) Optical mosaics (20 × 20 mm lattices, design pitch = 1,500 µm) printed at head speeds from 5 to 110 mm s^−^¹; right-hand panels show magnifications of foursquare pores. (B) DSC of PCL used for NF-MEW printing to find the melting and crystallization point (T^−^ = 89 °C; print temperature = 100 °C). Evolution of median pore width with speed; values plateau within ± 5 % of the CAD pitch once speed exceeds 30 mm s^−^¹. Box-and-whisker plots of strand diameter for the first, mid, and last strands at each speed (n = 10); asterisks denote Tukey post-hoc significance (**** p < 0.0001, *** p < 0.001, ** p < 0.01, ns = not significant). (E1) Detail views of four candidate speeds (50, 60, 70, 100 mm s^−^¹) printed at the optimized pressure of 3 kPa; dashed ellipses highlight residual waviness (60 mm s^−^¹) and discontinuous tracks (100 mm s^−^¹). (E2) Mean ± SD strand diameters for the same four speeds and comparison of fibre diameter on the nonwoven-PCL back-side versus MEW front-side with Tukey significance indicated. (E3) Pore-width distribution across the four candidate speeds, all within the 1,495–1,505 µm acceptance band (NS = no significant difference). Collectively, the data identify 70 mm s^−^¹ as the speed that yields straight, continuous strands, the lowest vertical coefficient of variation, and pore dimensions that match the CAD design while shortening build time by 14 % relative to the 60 mm s^−^¹ baseline.

This qualitative trend was quantitatively corroborated by pore size analysis (Fig. 2C). The pore size data provides a quantitative measure of printing fidelity. At low speeds (5-10 mm/s), the average pore size was small (∼800-1100 µm) and highly variable, a direct result of pore occlusion from fiber coiling. As the speed increased through the transitional regime, the average pore size steadily grew. Crucially, upon entering the stable regime (80-110 mm/s), the average pore size plateaued at a consistent value of ∼1450 µm with significantly reduced variability. The observed stabilization of the experimental architecture clearly demonstrates its convergence with the theoretical design, thereby signifying a high-fidelity printing process and confirming its success. Table 2 provides a comprehensive overview of the printing parameters that correspond to each distinct printing speed, offering a detailed breakdown for each setting.

**Table 2:** Printability metrics with the relative build-time recalculated against a 60 mm s^−^¹ baseline.

| Speed<br>(mm s<sup>-1</sup>) | Strand Ø –<br>1st (µm) | Strand Ø – mid<br>(µm) | Strand Ø –<br>last (µm) | CV<sub>Vert</sub> (%) † | Pore<br>width<br>(µm) | Relative print<br>time‡ |
| --- | --- | --- | --- | --- | --- | --- |
| 5 | 50.2±6.2 | 26.9±4.3 | 28.5±4.3 | 30 | 797±104 | 12.0 × baseline |
| 10 | 47.5±5 | 31.4±1.6 | 29.3±2.1 | 23 | 1067±75 | 6.0× |
| 20 | 33.5±3.3 | 19.8±2.4 | 20.1±2.8 | 26 | 1314±49 | 3.0× |
| 30 | 38.0±3.0 | 21.8±2.1 | 19.8±1.6 | 31 | 1339±48 | 2.0× |
| 40 | 33.9±2.6 | 20.6±1.8 | 19.8±1.06 | 26 | 1372±70 | 1.50× |
| 50 | 26.0±4.4 | 23.9±2.7 | 16.2±1.8 | 19 | 1442±83 | 1.20× |
| 60 | 31.9±5.3 | 24.4±3.4 | 22.5±2.1 | 15 | 1436±70 | 1.00× (baseline) |
| 70 | 46.5±2.1 | 34.1±4.4 | 31.3±1.8 | 18 | 1476±68 | 0.29× |
| 80 | 37.9±4.8 | 28.8±3.7 | 25.4±2.9 | 17 | 1442±44 | 0.25× |
| 90 | 40.4±3.26 | 29.5±2.7 | 27.7±1.4 | 17 | 1457±40 | 0.22× |
| 100 | 43.8±4.4 | 29.5±2.9 | 27.3±2.3 | 22 | 1459±33 | 0.20× |
| 110 | 42.2±3.8 | 28.7±2.3 | 29.0±2.4 | 19 | 1455±44 | 0.18× |

† Coefficient of variation of the three strand means (first, mid, last) at each speed.
‡ Relative build-time factor is calculated as 60/V; values < 1 indicate a faster print than the 60 mm s<sup>-1</sup> baseline, values > 1 a slower print.

Relative print-time expresses how long a build takes compared with a reference speed (*V_ref_*). Assuming the tool-path length (*L*) is identical for every specimen and the stage moves at constant velocity, build time is: 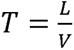

Normalizing to the reference run gives:

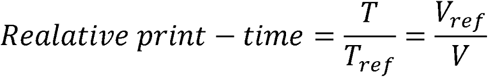

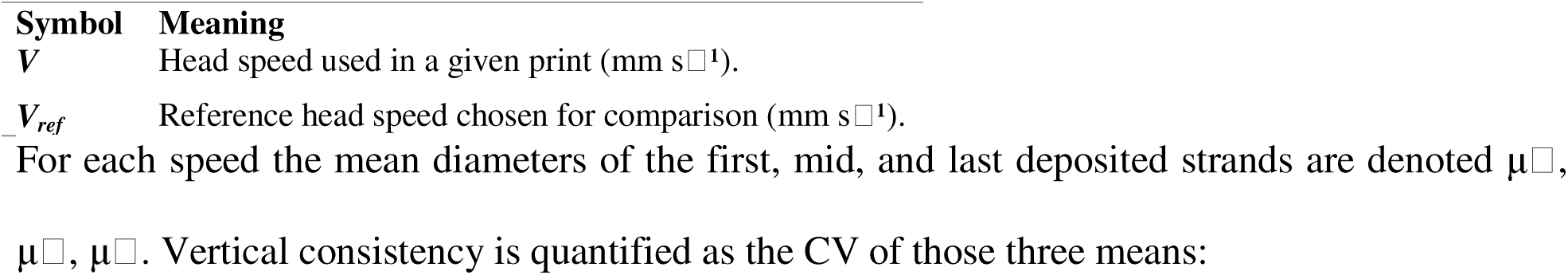

For each speed the mean diameters of the first, mid, and last deposited strands are denoted μ_1,2,3_,

μ_1,2,3_, μ_1,2,3_. Vertical consistency is quantified as the CV of those three means:

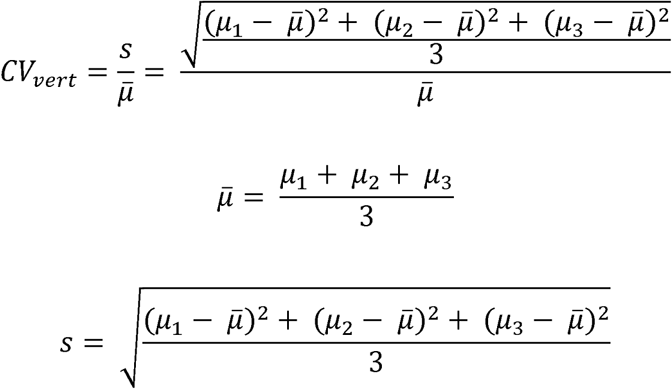

| Symbol | Meaning |
| --- | --- |
| $\mu_1$ | Mean diameter of the first deposited strand. |
| $\mu_2$ | Mean diameter of a mid-print strand (typically halfway through the build). |
| $\mu_3$ | Mean diameter of the last deposited strand. |
| $\bar{\mu}$ | Arithmetic mean of $\mu_1$, $\mu_2$, $\mu_3$ (overall average strand diameter). |
| $s$ | Standard deviation of the three strand means (measure of vertical scatter). |
| $CV_{vert}$ | Coefficient of variation; quantifies vertical uniformity (dimensionless). |

A lower CV*_vert_*indicates a more uniform strand diameter from the first to the last deposition pass.

Further investigation into intra-print consistency at 5 kPa revealed key insights into the process dynamics (Fig. 2D). A consistent “first strand effect” was observed, where the first deposited strand was significantly thicker than those printed subsequently. However, within the high-speed stable window (80-110 mm/s), the process achieved a steady state, with no significant difference in diameter between the middle and last strands. This suggested that while high speeds promoted consistency during the bulk of the print, the process could be further refined to improve initiation and overall robustness.

Based on the initial findings, a refinement study was conducted by reducing the printing pressure to 3 kPa, aiming to improve process control. Four candidate print-head speeds (50, 60, 70, and 100 mm/s¹) were evaluated for strand fidelity and vertical uniformity (Fig. 2E_1_–E_3_).

At 50 mm s^−^¹, large loops were suppressed, yet many horizontal rasters still adopted a shallow sinusoidal profile, whereas vertical rasters remained rectilinear (Fig. 2E_1_). The anisotropy arises because the jet briefly falls below its critical translation speed (CTS) after 90° rotation, allowing a compressive buckling wave to freeze in the X-direction. Statistically, the first strand was almost twice as thick as the mid strand and the last strand rebounded significantly (p < 0.001 for First vs Mid and Last vs Mid), yielding the poorest vertical-consistency coefficient (CV*_vert_* = 44%).

At 60 mm s^−^¹, raising the speed reduced diameter drift (CV*_vert_* = 39%) and removed any difference between mid and last strands (p = 0.69). Nevertheless, optical micrographs still showed recurrent wavy and bent segments, indicating residual jet-lag relative to the programmed tool-path (Fig. 2E_1_).

At 70mm s^−^¹, all filaments were straight, continuous and well registered. Mid and last strands were statistically identical (p = 0.73) and the first-strand overshoot dropped to a modest level (p = 0.012), giving CV*_vert_* = 24% —the lowest among defect-free builds—while shortening build time to 0.86 × the 60 mm s^−^¹ baseline. This speed therefore balances accuracy, consistency, and throughput (Fig. 2E_1_).

At 100 mm s^−^¹, although the three strand means were statistically indistinguishable (p > 0.42; CV*_vert_* = 2%), multiple broken or discontinuous tracks appeared (Fig. 2E_1_), proving that 3 kPa cannot supply melt fast enough at this traverse speed. The case underscores that numeric consistency alone can hide catastrophic defects and must be corroborated visually.

In stark contrast, the 70 mm/s speed emerged as the unequivocally optimized parameter. It successfully balanced all requirements, producing straight, continuous fibers free of defects and demonstrating excellent intra-print consistency. Notably, the significance level between the first strand and subsequent strands was reduced (p < ***), and the middle and last strands were statistically identical (ns), confirming a highly stable, steady-state process.

Taken together, these observations confirm 70 mm s^−^¹ as the robust operating point for subsequent scaffold fabrication: it eliminates sinusoidal artefacts, maintains strand-to-strand uniformity, and offers a meaningful gain in throughput without risking flow interruption.

The final confirmation came from pore size analysis under the refined conditions (Fig. 2E_3_). While the pore sizes were not statistically different across the tested groups, only the 70 mm/s speed produced scaffolds with a median pore size that precisely matched the intended architectural threshold. The other speeds resulted in pores that were measurably below this target (Table 3).

**Table 3:** Mean ± SD strand diameters, vertical-consistency coefficient of variation (CV_vert_) and relative build-time (normalized to 60 mm s^−^¹) for meshes printed at 3 kPa. CV_vert_ is calculated from the first, mid, and last strand means; lower values denote greater uniformity through the build.

| Speed<br>(mm s<sup>-1</sup>) | Strand Ø – 1st<br>(µm) | Strand Ø – mid<br>(µm) | Strand Ø – last<br>(µm) | $CV_{vert}$*<br>(%) | Pore width<br>(µm) | Relative<br>time† |
| --- | --- | --- | --- | --- | --- | --- |
| 50 | 34.64±3.02 | 14.58±2.29 | 15.03±3.04 | 44 | 1490.16±16.63 | 1.20× |
| 60 | 25.87±1.50 | 13.20±2.17 | 11.22±1.75 | 39 | 1493.28±13.23 | 1.00× |
| 70 | 10.89±2.09 | 7.33±2.32 | 6.41±1.60 | 24 | 1498.95±6.93 | 0.86× |
| 100 | 21.73±1.33 | 22.95±7.40 | 22.66±2.37 | 2 | 1491.57±8.09 | 0.60× |

\* $CV_{vert}$ = standard deviation of the three strand means ÷ their grand mean.
†Relative print-time = 60 / speed; values $< 1$ indicate a faster build than the 60 mm s<sup>-1</sup> reference.

Through a systematic, two-stage optimization, the ideal processing parameters for fabricating high-fidelity PCL scaffolds were determined by precisely managing the dynamic relationship between the polymer jet’s intrinsic velocity (V*_jet_*) and the critical translation speed (V*_CTS_*).

The initial screening at 5 kPa pressure clearly illustrated this principle. At low speeds, where V*_jet_* was greater than V*_CTS_* (V*_jet_* > V*_CTS_*), the excess extruded material resulted in fiber coiling and instability.

A stable process, characterized by straight fibers, was achieved at high speeds (80-110 mm/s), establishing a condition where V*_CTS_* was greater than V*_jet_*(V*_CTS_* > V*_jet_*), enabling stabilizing mechanical drawing of the jet.

Following the initial process, a subsequent refinement step, involving a reduction of pressure to 3 kPa, resulted in a shift of the optimal operational window. At 60 mm/s, the re-emergence of wavy fibers suggests the system was operating too close to a balanced condition (V*_jet_* ≈ V*_CTS_*), where minor perturbations could still cause instability.

At 100 mm/s, the observation of fiber breakage indicates a condition where V*_CTS_* was much greater than V*_jet_*(V*_CTS_* >> V*_jet_*). Here, the drawing forces exceeded the tensile limit of the molten polymer jet, causing it to fracture before solidification.

Ultimately, the optimized combination of 3 kPa pressure and 70 mm/s V*_CTS_* was identified as the ideal processing state. This parameter set establishes the most favorable V*_CTS_* > V*_jet_* condition: it is fast enough to guarantee stabilizing jet drawing for exceptional fidelity and consistency, yet remains below the critical speed that would cause material failure. This robust process, which balances these competing forces, is ideally suited for fabricating geometrically complex and highly reproducible scaffolds.

### 3.2. Barrier Film Assembly

Following the optimization of the MEW process, a multi-layered, bio-composite construct designed to function as a uterine barrier film was successfully assembled. The design integrates a structural PCL framework with a functional hydrogel derived from decellularized uterine extracellular matrix (dUECM) and a cellular layer, as illustrated schematically and confirmed with SEM analysis in Figure 3.

**Figure 3:**
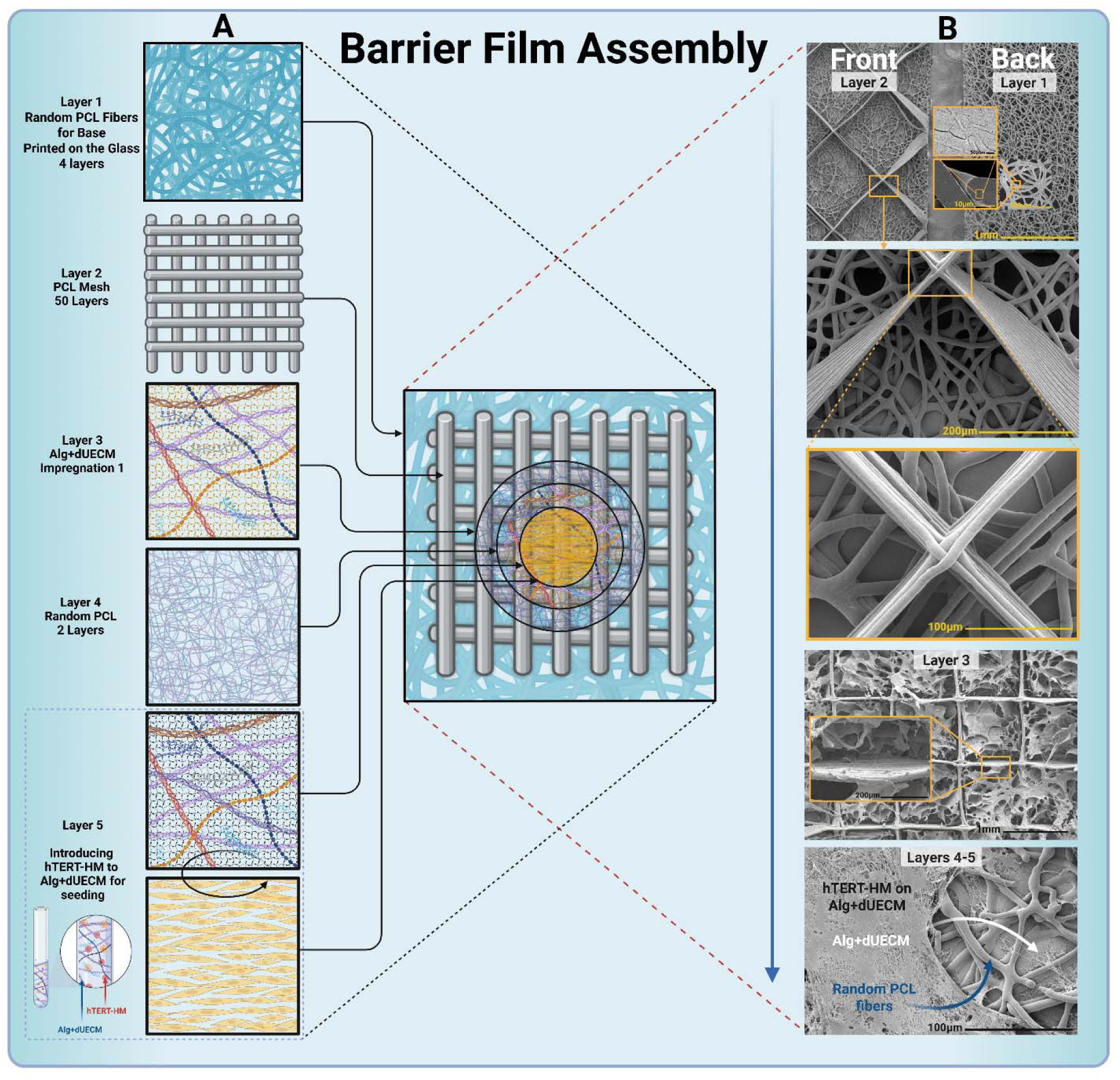
Multilayer barrier-film design and morphology. Panel (A) shows a layer-by-layer fabrication sequence: nonwoven PCL base (4 layers) → MEW mesh (50 layers) → Alg+dUECM impregnation → nonwoven PCL cover (2 layers) → Alg+dUECM/hTERT-HM seeding. Panel (B) The right panel shows SEM micrographs of the front (mesh side) and back (nonwoven-fibre side) interfaces; the inset shows a fused cross-over junction. Layer 3 shows the Alg+dUECM-filled mesh pores, followed by the top view after 48 h of culture, showing hTERT-HM cells spanning the hydrogel and underlying nonwoven PCL fibers.

The construct was designed as a multi-laminar assembly with distinct functional layers (Fig. 3A). To begin the procedure, Layer 1 was implemented as a base layer, incorporating four nonwoven oriented layers of PCL fiber to encourage hydrophobic side behavior and initial anti-adhesion properties. This was followed by 50 layers of highly ordered PCL mesh, fabricated using the optimized MEW parameters, intended to provide primary structural integrity and defined porosity to support the bioactive hydrogel (Layer 2). Subsequently, the PCL meshes (90°, 60°, 45°, and 30°) were impregnated with a hydrogel composite of Alg+dUECM (Layer 3). A thin layer of nonwoven PCL fibers (Layer 4) was printed over freeze-dried Mesh/Alg+dUECM hydrogel to enhance mechanical cohesion and provide a scaffold for the final cell layer. The assembly was completed by seeding hTERT-HM cells mixed with Alg+dUECM hydrogel as a carrier onto the surface (Layer 5), creating the final functional barrier film.

The SEM images clearly distinguish the highly ordered PCL mesh of Layer 2 on the “Front” from the underlying, fine, nonwoven fiber mat of Layer 1 on the “Back” (Fig. 3B). Magnified views of the MEW junctions show clean, fused fiber contacts, demonstrating excellent intra-layer structural integrity.

The image corresponding to Layer 3 shows that the Alg+dUECM hydrogel has been successfully impregnated into the pores of the PCL mesh. The hydrogel forms a continuous phase that fills the micro-sized pores, effectively creating a hydrophilic barrier within the structural framework. The cross-sectional inset confirms that the hydrogel completely embeds the PCL fibers, creating a true composite structure.

The final surface morphology shows a complex but well-integrated system. The larger, nonwoven PCL fibers from Layer 4 are visible, providing a textured substrate. These fibers are coated and embedded within the second hydrogel impregnation (Layer 5). Notably, the hTERT-HM cells are observed adhering and spreading across this composite surface, forming a confluent monolayer, as indicated by the arrows. The final construct has been confirmed as successfully supporting both cell attachment and cell proliferation, indicating its potential for use in further experiments.

### 3.3. Thermogravimetric Analysis (TGA) of PCL and PCL/Alg+dUECM composite

To rigorously investigate the thermal stability and verify the successful integration of all components into the final construct, Thermogravimetric Analysis (TGA) and its first derivative (DTG) were conducted. The results, comparing the pure PCL mesh to the final PCL Mesh / Alg-dUECM composite, are presented in Figure 4A (TGA) and Figure 4B (DTG).

**Figure 4:**
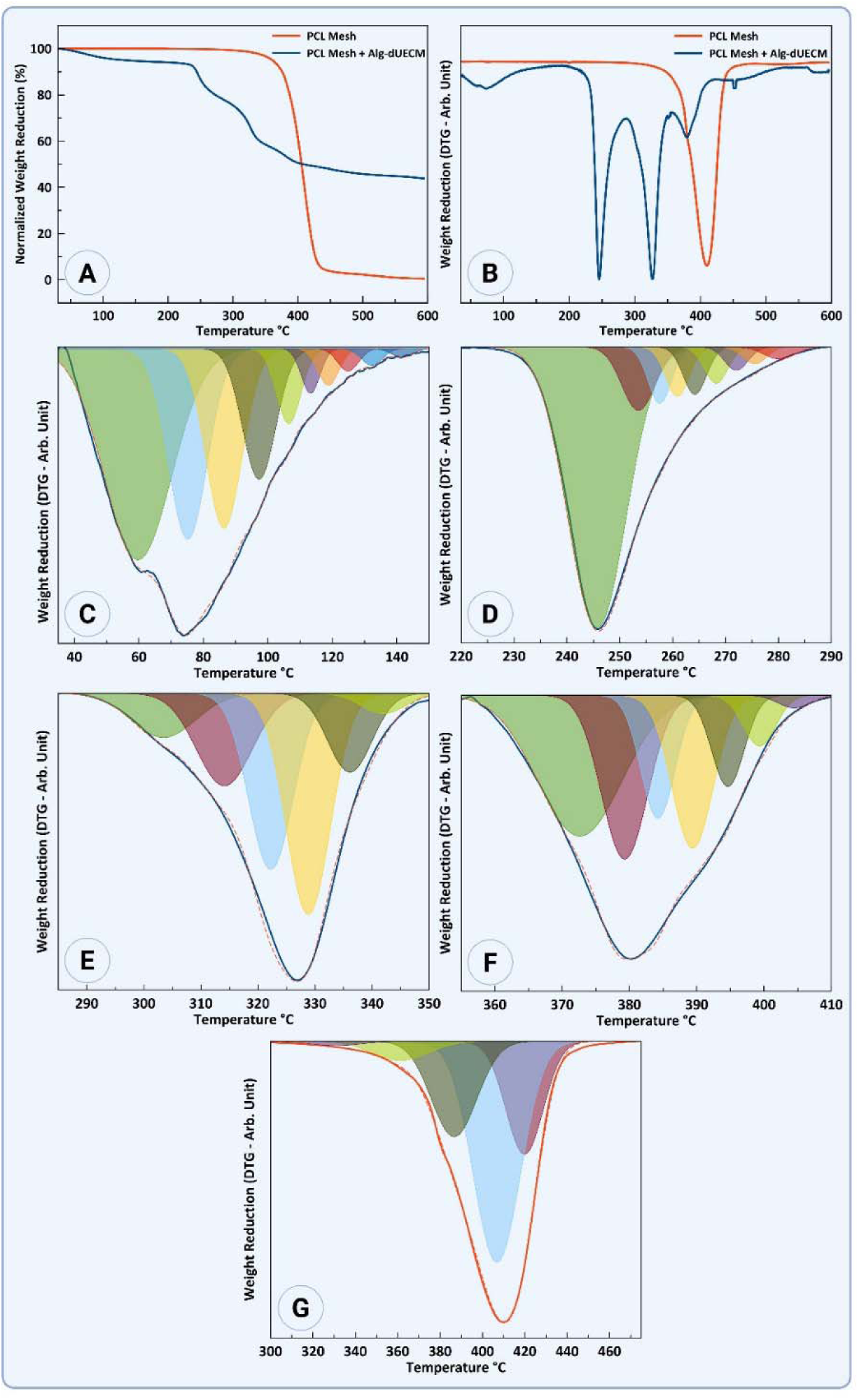
Thermogravimetric Analysis (TGA) confirming the successful fabrication and multi-component nature of the PCL/Alg-dUECM composite film. (A) TGA thermograms showing the normalized weight reduction as a function of temperature for the pure PCL Mesh (red) and the PCL Mesh + Alg-dUECM composite (blue). The pure PCL displays a single, sharp degradation step, while the composite exhibits a clear multi-stage degradation profile. (B) Corresponding first derivative thermogravimetry (DTG) curves, highlighting the rate of weight reduction. The PCL mesh shows a single peak, whereas the composite shows multiple distinct peaks indicative of its different components. (C-G) Magnified views of the DTG peaks for the composite film, pinpointing specific thermal events: (C) Initial weight loss below 150 °C corresponding to water evaporation from the hydrogel components. (D) A sharp peak at ∼245 °C attributed to the decomposition of the alginate backbone. (E, F) A broad, multi-peak region between 290 °C and 400 °C representing the complex degradation of various proteins within the dUECM. (G) The final degradation peak above 400 °C, present in both samples, corresponding to the thermal breakdown of the PCL polymer backbone.

The PCL mesh only (red line) exhibits the classic thermal profile of a high-molecular-weight, semi-crystalline polymer. It demonstrates high thermal stability with a single, well-defined degradation step. The onset of thermal decomposition begins at approximately 350 °C, followed by a rapid and near-complete mass loss, which concludes by 450 °C, leaving negligible residual char. The corresponding DTG curve shows a single, sharp peak with a maximum rate of degradation (T*_peak_*) centered at ∼415 °C (Figure 4G). This degradation event corresponds to the nonwoven chain scission of the ester linkages in the PCL backbone at elevated temperatures [27].

A complex, multi-stage degradation profile, clearly visible in the composite film (blue line), stands in sharp contrast to simpler profiles seen elsewhere, providing undeniable evidence of the material’s heterogeneous and complex internal composition. This profile can be deconstructed into four key thermal events, each corresponding to a specific component within the construct:

1. **Initial Dehydration:** A significant weight loss of approximately 10% is observed in the low-temperature region between 40 °C and 150 °C. This initial mass reduction, which is entirely absent in the hydrophobic PCL sample, is attributed to the evaporation of free and bound water molecules physically entrapped within the hydrophilic hydrogel network of alginate and dUECM components (Fig. 4C).
2. **Alginate Depolymerization:** Following dehydration, the first major structural decomposition occurs with a sharp and prominent DTG peak centered at ∼245 °C. This peak corresponds to the thermal degradation of sodium alginate, which involves the dehydration of saccharide rings and the cleavage of glycosidic bonds (Fig. 4D).
3. **dUECM Protein Denaturation and Decomposition:** The temperature range from 290 °C to 400 °C is characterized by a broad and complex degradation pattern with multiple overlapping peaks, most notably at ∼325 °C and ∼380 °C. This signature is indicative of the decomposition of the protein-rich dUECM. Biological matrices are composed of a diverse array of proteins (e.g., collagen, elastin, glycoproteins), each with a distinct thermal stability, resulting in a multi-faceted degradation profile rather than a single sharp peak (Fig. 4E and F).
4. **PCL Backbone Degradation within the Composite:** The final degradation stage, with a T*_peak_* at ∼420 °C, clearly corresponds to the thermal breakdown of the PCL component. Notably, the degradation peak of PCL within the composite is shifted to a slightly higher temperature compared to the pure PCL mesh (∼415 °C). This phenomenon suggests a mild “protective effect,” where the char formed from the decomposition of the alginate and dUECM components at lower temperatures may act as a temporary thermal barrier, slightly inhibiting the degradation of the PCL framework (Fig. 4G).

### 3.4. Chemical Composition and Functional Group Analysis by FTIR Spectroscopy

Fourier Transform Infrared (FTIR) spectroscopy was performed to confirm the chemical identity of the PCL mesh and to provide definitive proof of the successful integration of the Alg-dUECM hydrogel into the final composite film. Figure 5A presents the full spectra, while Figures 5B1-C3 show detailed, deconvoluted analyses of specific spectral regions.

**Figure 5:**
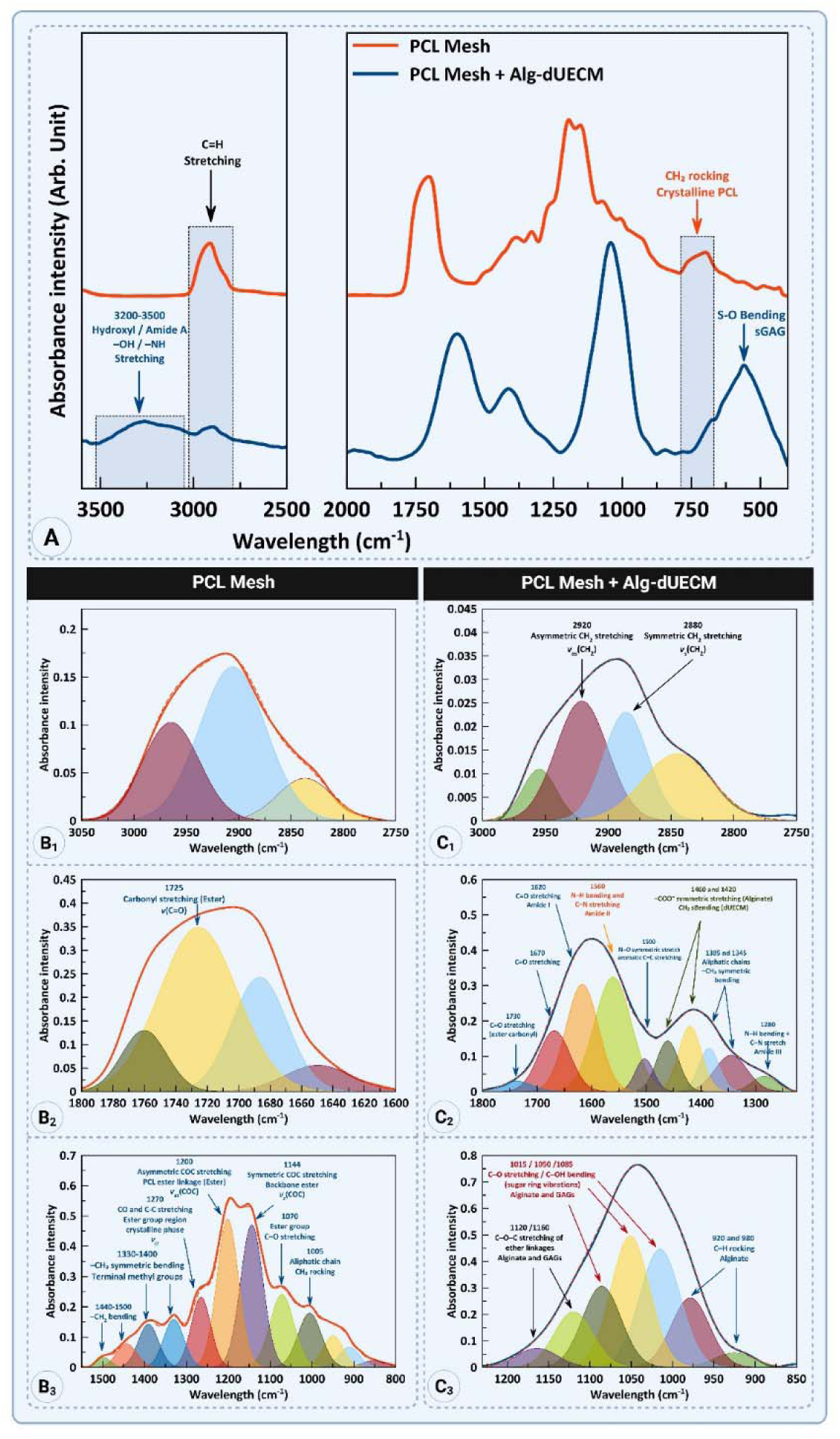
Chemical verification of PCL/Alg-dUECM composite formation by FTIR spectroscopy, confirming the successful molecular integration of the hydrogel components within the PCL mesh. (A) Full-range FTIR spectra comparing the PCL Mesh (red) with the final PCL Mesh + Alg-dUECM composite (blue). Note the prominent emergence of a broad hydroxyl/amide band (3200-3500 cm^−^¹) and significant modifications in the fingerprint region (1700-500 cm^−^¹) in the composite spectrum, indicating the presence of new functional groups. (B1-B3) Detailed analysis of key spectral regions for the pure PCL mesh, showing its characteristic chemical signature: (B1) C-H stretching vibrations; (B2) the dominant ester carbonyl (C=O) peak at 1725 cm^−^¹; and (B3) the C-O-C stretching and CH^−^ bending peaks in the fingerprint region. (C1-C3) Detailed analysis of the PCL Mesh + Alg-dUECM composite, highlighting the new peaks that confirm successful integration: (C1) The C-H region, now broadened by the presence of -OH and -NH groups. (C2) Definitive evidence of the dUECM, showing the protein-specific Amide I (∼1650 cm^−^¹) and Amide II (∼1560 cm^−^¹) bands, alongside the carboxylate (–COO^−^) peaks (1460 and 1420 cm^−^¹) from alginate. (C3) The emergence of a complex and intense polysaccharide band (1015-1160 cm^−^¹) from C-O and ether linkage vibrations in the alginate and GAGs.

The FTIR spectrum of the pure PCL mesh (red line) exhibits all the characteristic absorption bands of Polycaprolactone. The C-H stretching region (Fig. 5B1) shows two prominent peaks corresponding to the asymmetric (ν*_as_*(CH )) at 2940 cm^−^¹ and symmetric (ν*_s_*(CH )) C-H stretching at 2865 cm^−^¹ of the methylene groups. The most dominant feature of the spectrum (Fig. 5B2) is the intense, sharp absorption peak at 1725 cm^−^¹, which is the signature stretching vibration of the carbonyl group (ν(C=O)) within the ester linkage of PCL. The fingerprint region (Fig. B3) displays several characteristic PCL peaks, including the asymmetric C-O-C stretching at 1293 cm^−^¹ and symmetric C-O-C stretching at 1144 cm^−^¹ [28]. The absence of significant peaks in the 3200-3500 cm^−^¹ range confirms the lack of hydroxyl or amide groups in the pure polymer.

The spectrum of the PCL Mesh + Alg-dUECM composite (blue line) is markedly different, providing chemical confirmation of hydrogel integration by displaying characteristic bands of all its distinct components. The most striking difference is the appearance of a new, intense, broad absorption band between 3200 and 3500 cm^−^¹ (Fig. 5A). This band is a composite of overlapping O-H stretching vibrations from hydroxyl groups in alginate and glycosaminoglycans (GAGs), as well as N-H stretching vibrations (Amide A) from the protein components of the dUECM [29].

The carbonyl region (Fig. 5C2) is profoundly altered. While the PCL ester peak at ∼1730 cm^−^¹ is present, it is now accompanied by two strong peaks that are definitive markers for proteins: Amide I band (1620-1670 cm^−^¹), which is arising primarily from C=O stretching within the peptide bonds. Amide II band (∼1560 cm¹), which results from N-H bending and C-N stretching in the peptide backbone [30].

The composite spectrum exhibits key features that confirm the presence of polysaccharide components. Characteristic peaks for alginate’s carboxylate groups (–COO ) appear at 1460 and 1420 cm^−^¹ (Fig. 5C2) [30, 31]. A broad and intense region between 1015 and 1160 cm^−^¹ corresponds to complex C-O and C-O-C vibrations of the pyranose rings in both alginate and dUECM GAGs (Fig. 5C3). The peak labeled S-O Bending suggests the presence of sulfated GAGs (sGAGs), which are integral components of the native uterine ECM [4, 32].

In summary, the FTIR analysis provides unequivocal chemical evidence of composite fabrication. The presence of new, distinct absorption bands for hydroxyl/amide groups, Amide I and II bands, and carboxylate/saccharide vibrations, which are entirely absent in the pure PCL spectrum, definitively confirms the homogenous integration of the Alg-dUECM hydrogel throughout the PCL mesh framework.

### 3.5. Mechanical Properties of the PCL + dUECM MEW printed Scaffolds

To evaluate the mechanical suitability of the composite scaffolds for uterine repair, their tensile properties were comprehensively characterized and benchmarked against native uterine tissue. Recognizing that the material properties are highly dependent on their hydration state, tests were conducted under both dry (freeze-dried) and wet (physiologically relevant) conditions. The influence of the MEW fiber arrangement (90°, 60°, 45°, and 30°) on mechanical anisotropy was also investigated, both initially and after a 7-day incubation period in cell culture media to assess functional stability (Fig. 6).

**Figure 6:**
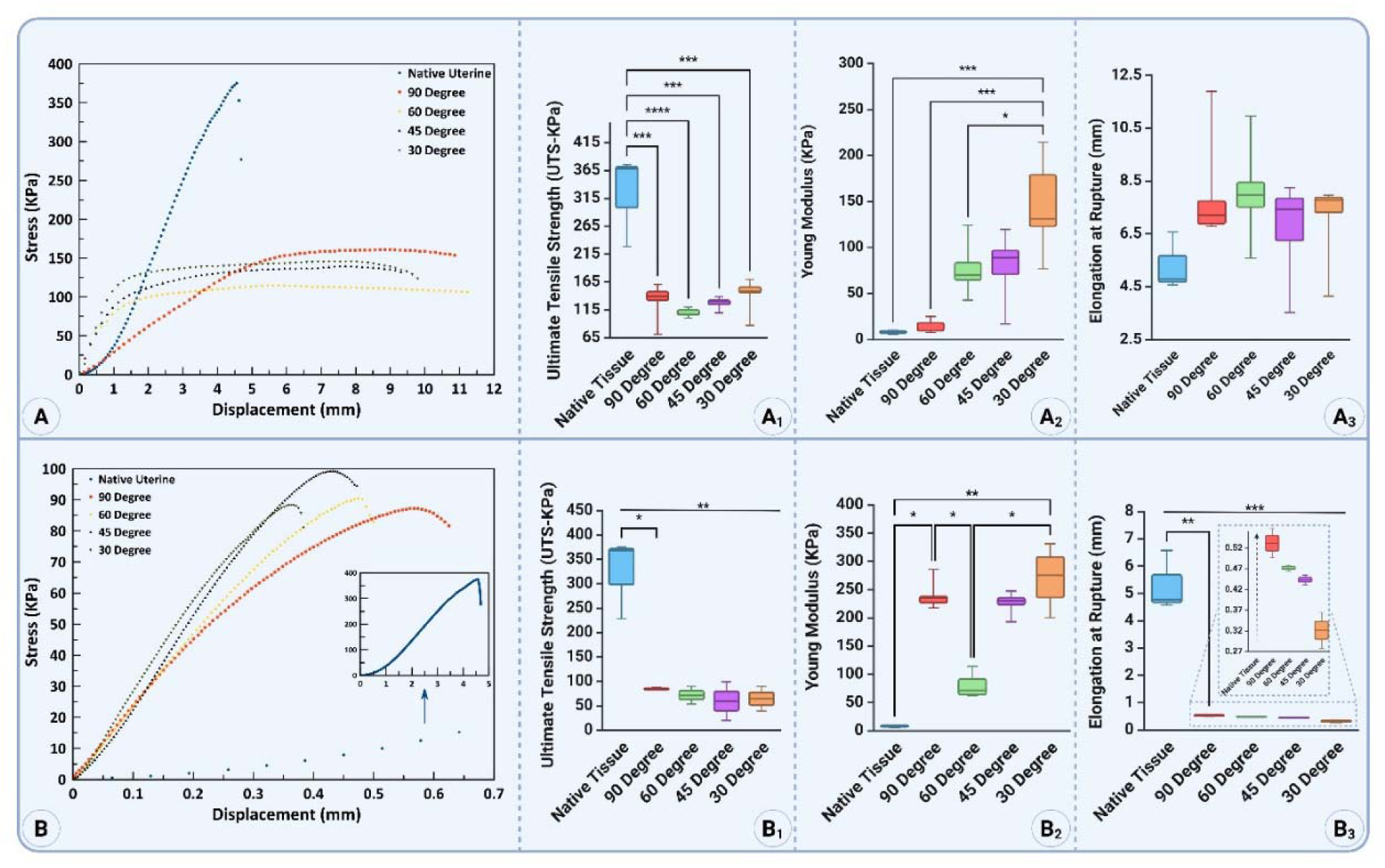
Mechanical characterization of the PCL/Alg-dUECM composite scaffolds under dry and wet conditions, benchmarked against native uterine tissue. (A) Representative stress-displacement curves for dry (freeze-dried) scaffolds with different angles compared to native uterine tissue. (A1-A3) Box plots summarizing the quantitative analysis of the dry samples for Ultimate Tensile Strength (UTS), Young’s Modulus, and Elongation at Rupture, respectively. (B) Representative stress-displacement curves for wet (hydrated) scaffolds, demonstrating the significant change in mechanical behavior upon hydration. (B1-B3) Box plots summarizing the UTS, Young’s Modulus, and Elongation at Rupture for the wet samples, respectively. The inset in (B3) provides a magnified view of the low elongation values characteristic of the engineered scaffold groups. All data are presented as mean ± SD. Statistical significance is denoted as *p < 0.05, **p < 0.01, ***p < 0.001, and ****p < 0.0001.

In the dry state, which represents a potential storage condition, the native uterine tissue proved to be the most robust material. The native uterine tissue exhibited a mean ultimate tensile strength (UTS) of 323±82 kPa, which was significantly stronger than all engineered scaffold groups (p < 0.0001). Tukey’s post-hoc analysis confirmed that the mean strength of native tissue was higher than the 90°, 60°, 45°, and 30° groups (130±34 kPa, 109±7 kPa, 127±12 kPa, and 141±31 kPa, respectively). There were no statistically significant differences in UTS observed between any of the scaffold designs themselves (all adjusted p-values > 0.6), indicating that fiber architecture did not alter the UTS in the dry state (Fig. 6A1).

Upon analysis of the scaffold’s stiffness, it was revealed that the Young’s modulus of the material could be effectively tuned via manipulation of its underlying architecture. The 30° design was significantly stiffer than the native tissue (p = 0.0004), as well as being significantly stiffer than both the 90° (p = 0.001) and no different significance with 60° and 45° designs (p = 0.095 and p = 0.1073, respectively). While a trend towards increased stiffness was observed for the 45° and 60° groups, these differences were not statistically significant (p = 0.999) (Fig. 6A2).

In terms of elongation, no statistically significant differences in elongation at rupture were observed between any of the groups (all adjusted p-values > 0.29), suggesting that all materials possess a similar capacity for stretching before failure in the dry condition (Fig. 6A3).

To assess the functional stability of the composite scaffolds over time, their mechanical properties were tested after a 7-day incubation period in DMEM/F12 cell culture media. This experiment simulates the performance of the acellular scaffold in a physiological environment prior to significant cell-mediated remodeling or degradation.

After 7 days, all engineered scaffold groups maintained substantial mechanical integrity. The 90° design exhibited the highest mean UTS at 84.6±3.6 kPa, followed by the 60° (71.7±25.4 kPa), 30° (64.4±35.7 kPa), and 45° (59.7±55.9 kPa) designs. Despite these numerical differences in the mean values, an analysis comparing the different architectural designs found no statistically significant differences in ultimate strength among the 90°, 60°, 45°, and 30° groups (Fig. 6B1). This indicates that while the fiber lay-down angle influences other properties, it does not significantly alter the material’s ultimate strength after a 7-day incubation in a hydrated state.

The stiffness of the scaffolds remained high after the 7-day incubation period, with architectural dependence being a key feature. Dunn’s post-hoc test showed that both the 90° and 30° scaffolds were significantly stiffer than the native tissue (p = 0.021 and p = 0.007, respectively). Furthermore, differences among the scaffold designs persisted, with the 30° scaffold being significantly stiffer than the 60° scaffold (p = 0.012) (Fig. 6B2). This confirms that architectural control of stiffness is a durable feature of the design.

The elongation of the different scaffold designs was found to be highly consistent, with all groups exhibiting very low stretchability before failure. The mean elongation values were 0.53 mm for the 90° design, 0.47 mm for the 60° design, 0.44 mm for the 45° design, and 0.32 mm for the 30° design (Fig. 6B3). Although a trend was observed showing a decrease in elongation with increasing acuteness of angles, a comprehensive post-hoc analysis using Tukey’s method revealed the absence of any statistically significant variations in elongation at rupture across the different engineered scaffold designs, indicating no discernible differences between the groups.

In summary, the 7-day incubation study demonstrates that composite scaffolds are mechanically stable in a simulated physiological environment. The key finding is that the stiffness (Young’s Modulus) can be durably and significantly tuned by altering the MEW fiber architecture. In contrast, the UTS and elongation remain consistent across the different designs in the functional, hydrated state.

### 3.6. Swelling and Degradation Profiles of the Composite Scaffolds

The ability of the composite scaffolds to absorb and retain media, a critical function of the integrated hydrogel component, was quantitatively assessed by measuring the swelling ratio over an 18-hour (1140-minute) period. The results, calculated from the provided data, are detailed below and presented in Figure 7A.

**Figure 7:**
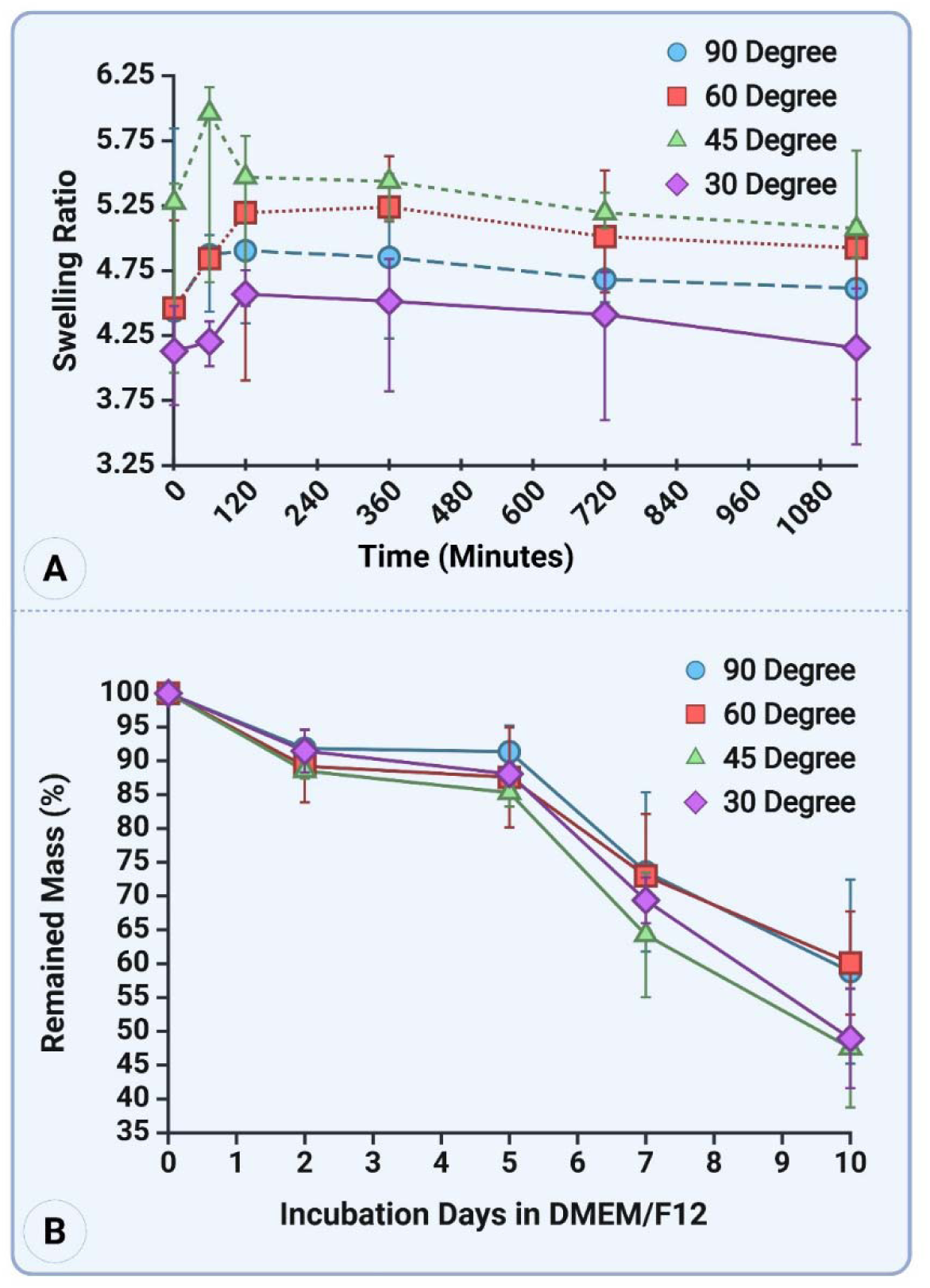
Functional characterization of the PCL/Alg-dUECM composite scaffolds, demonstrating architectural control over swelling kinetics and in vitro degradation. (A) Swelling ratio profiles of the scaffolds with different fiber angles measured over an 18-hour (1080-minute) period. All designs show rapid initial media uptake leading to an equilibrium swelling state, the magnitude of which is clearly dependent on the scaffold’s architecture. (B) In vitro degradation analysis showing the percentage of remaining scaffold mass over a 10-day incubation period in DMEM/F12 cell culture media. The results demonstrate a controlled, progressive mass loss, with the rate of degradation also being influenced by the architectural design. All data points are presented as mean ± standard deviation (SD).

All composite scaffolds exhibited the characteristic behavior of a hydrogel-containing material. A phase of rapid initial media uptake occurred within the first 60-120 minutes, demonstrating the immediate hydrophilic nature conferred by the Alg-dUECM component. After this initial surge, the swelling rate decreased as the scaffolds gradually approached an equilibrium state.

The scaffold architecture was found to have a clear and significant influence on both the rate of swelling and the final equilibrium swelling ratio.

The 45° design consistently demonstrated the highest water absorption capacity. It reached a peak mean swelling ratio of 5.59 ± 0.84 within the first 60 minutes and stabilized at an equilibrium swelling ratio of 5.20 ± 0.42 by the 1140-minute time point, the highest among all groups. The 90° and 60° designs showed intermediate swelling behavior. The 90° design reached an equilibrium ratio of 4.68 ± 0.17, while the 60° design stabilized at a very similar value of 4.60 ± 0.77. The 30° design consistently exhibited the lowest swelling capacity. After an initial uptake, it equilibrated at a final swelling ratio of 4.06 ± 0.61, significantly lower than the other configurations.

The stability and degradation kinetics of the composite scaffolds were assessed over a 10-day incubation period in DMEM/F12 media to simulate their behavior in a physiological environment. The percentage of remaining mass was measured at key time points, with the results presented in Figure 7B.

All scaffold groups exhibited a progressive loss of mass throughout the 10-day study. This mass loss is primarily attributed to the degradation and dissolution of the Alg-dUECM hydrogel component, as the PCL framework degrades over a much longer timescale. The degradation profiles indicate that the architectural design of the MEW mesh has a significant influence on the rate at which the scaffold breaks down (Table 4).

**Table 4:** The DMEM/F-12 degradation confirms a biphasic profile for all raster angles (mean ± SD, n = 3 per time point)

| Day | 30° | 45° | 60° | 90° |
| --- | --- | --- | --- | --- |
| 0 | 100 % | 100 % | 100 % | 100 % |
| 2 | 91.5 ± 4.8 % | 88.5 ± 3.0 % | 89.2 ± 2.5 % | 91.9 ± 2.3 % |
| 5 | 88.1 ± 1.9 % | 85.3 ± 1.7 % | 87.6 ± 3.4 % | 91.3 ± 1.2 % |
| 7 | 69.4 ± 6.4 % | 64.3 ± 5.0 % | 73.0 ± 2.3 % | 73.6 ± 3.3 % |
| 10 | 48.9 ± 6.0 % | 47.5 ± 9.3 % | 60.1 ± 5.4 % | 58.8 ± 8.1 % |

By Day 5, all scaffolds showed a modest but clear mass loss. The 90° and 60° designs were most stable, retaining 91.3% ± 3.8% and 87.6% ± 7.4% of their initial mass, respectively. The more acute-angled scaffolds degraded slightly faster, with the 45° and 30° groups retaining 85.3% ± 2.1% and 88.1% ± 1.5% of their mass. By Day 7, the degradation rates had accelerated. The 90° design remained the most stable, retaining 73.6% ± 11.6% of its mass. The 60° design was slightly less stable at 73.0% ± 8.8%. Degradation was more pronounced in the 45° (64.3% ± 9.1%) and 30° (69.4% ± 3.4%) designs. By Day 10, significant mass loss was observed across all groups, and the architectural dependence became most apparent. The 90° design demonstrated the greatest stability, retaining 58.8% ± 13.6% of its mass. This was closely followed by the 60° design at 60.1% ± 7.7%. The 45° and 30° designs showed the most rapid degradation, retaining only 47.5% ± 8.8% and 48.9% ± 7.4% of their initial mass, respectively.

### 3.7. Surface Wettability and Hydrophilicity of Composite Scaffolds

The surface wettability of the scaffolds, a critical property for modulating protein adsorption and cell interaction, was systematically evaluated using contact angle goniometry with pre-warmed cell culture media (DMEM/F12). The analysis was designed to characterize the different functional surfaces of the construct and benchmark them against the corresponding native uterine tissue layers (Fig. 8).

**Figure 8:**
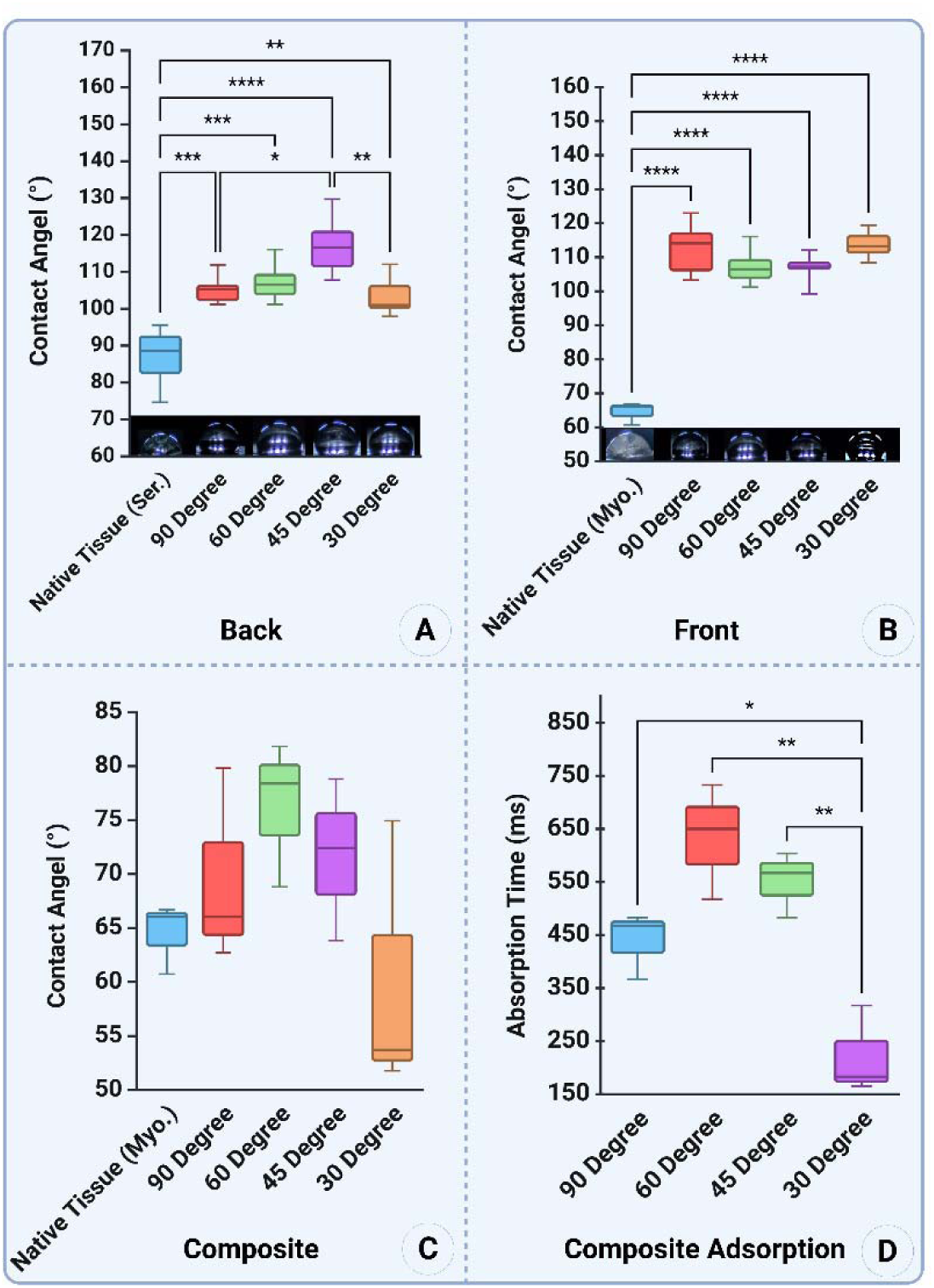
Surface wettability and absorption dynamics, demonstrating the successful functionalization of the scaffolds with a hydrophilic hydrogel. (A) Contact angle measurements on the hydrophobic “back” side (Layer 1, nonwoven PCL fibers) of the scaffolds, compared with the native tissue serosal layer. All PCL surfaces show significantly higher hydrophobicity than the native tissue. (B) Contact angle measurements on the hydrophobic “front” side (Layer 2, ordered MEW mesh) of the scaffolds prior to hydrogel impregnation, compared with the native tissue myometrial layer. All PCL architectures are confirmed to be highly non-wettable. (C) Contact angle measurements on the “front” side of the final composite scaffolds after impregnation with Alg-dUECM hydrogel. Note the dramatic decrease in contact angle for all scaffold designs, which now exhibit hydrophilic properties statistically similar to the native myometrium. (D) Quantitative analysis of the droplet absorption time for the hydrophilic composite scaffolds, showing architecture-dependent differences in the rate of fluid uptake. All data are presented as box-and-whisker plots showing median, interquartile range, and min/max values. Statistical significance is denoted as *p < 0.05, **p < 0.01, ***p < 0.001, and ****p < 0.0001.

First, the “back” surface of the construct (Layer 1, nonwoven PCL fibers) was analyzed (Fig. 8A). The PCL scaffold surface was found to be distinctly hydrophobic, exhibiting high contact angles that varied with fiber architecture. The 45° design showed the highest hydrophobicity with a mean contact angle of 117.1°, which was significantly greater than the 90° (105.3°, p < ***), 30° (103.3°, p < ****), and native tissue serosa (86.9°, p < ***) groups. The native tissue serosa itself was moderately hydrophobic but still significantly more wettable than all scaffold designs.

Next, the “front” side of the scaffolds, composed of the ordered MEW mesh but without the hydrogel impregnation, was evaluated against the native myometrium tissue (Fig. 8B). A similar trend was observed: the native myometrium was hydrophilic with a mean contact angle of 64.5°, whereas all pure PCL mesh architectures were highly hydrophobic, with mean contact angles consistently above 106°. This confirms that the PCL polymer itself is inherently non-wettable, a property that must be overcome for biological applications.

The most critical test involved analyzing the front side of the final composite scaffold after impregnation with the Alg-dUECM hydrogel (Fig. 8C). The results show a dramatic and complete reversal of surface properties.

Upon impregnation with the hydrogel, all scaffolds became strongly hydrophilic, and their wettability closely mirrored that of the hydrophilic native myometrium control (mean contact angle 64.5±3.2°). There were no statistically significant differences in contact angle between the native myometrium and the 90° (69.5±9.0°), 60° (76.3±6.7°), or 45° (71.6±7.5°) hydrogel-impregnated scaffolds. The 30° design became even more hydrophilic than the native tissue, with a median contact angle of just 60.1±12.8°.

The water droplet’s absorption into the scaffold was rapid because the composite surface exhibited a remarkably hydrophilic nature. The time required for full absorption was therefore used as an additional quantitative measure of surface interactivity (Fig. 8D).

The absorption time was found to be dependent on the scaffold architecture. The 30° scaffold exhibited the fastest absorption, with a mean time of only 221±83 ms. The 90° and 45° designs absorbed the droplet at intermediate rates of 439±62 ms and 551±61 ms, respectively. The 60° scaffold was the slowest to absorb the droplet, with a mean time of 633±108 ms. Statistically, the absorption time for the 60° design was significantly longer than for the 30° (p < *) and 45° (p < **) designs.

### 3.8. Biological Consequence of PCL Hydrophobicity: An Imperative for Surface Modification

Having established through FTIR, TGA, and wettability tests that the neat PCL mesh is an inherently hydrophobic and chemically distinct material, a critical experiment was conducted to validate the functional necessity of surface modification. Due to the poor wettability of the PCL scaffolds, our hypothesis was that the hTERT-HM cells would not attach, spread, or form a monolayer effectively, thereby preventing the formation of a functional barrier film necessary for the bioactive hydrogel’s synergistic action.

To test this, unmodified PCL scaffolds of varying architectures were co-cultured with hTERT-HM cells, and the resulting cell-scaffold interactions were analyzed by SEM (Fig. 9).

**Figure 9:**
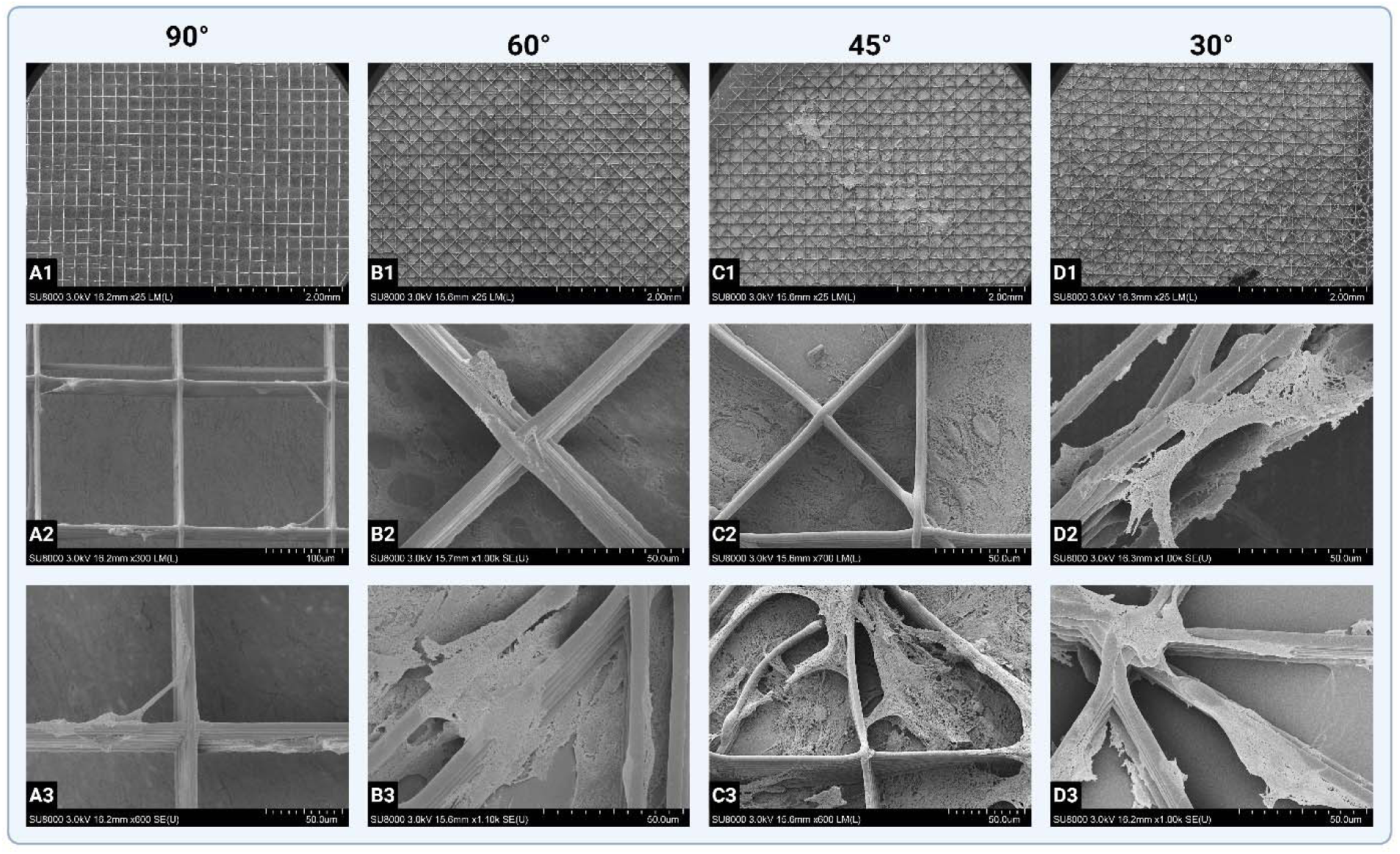
SEM micrographs demonstrating limited and selective hTERT-HM cell attachment to unmodified, hydrophobic PCL scaffolds, highlighting the need for surface functionalization. (A1-D1) Low-magnification overviews of the different scaffold architectures (90°, 60°, 45°, and 30°) after cell culture. (A2-A3) High-magnification views of the 90° scaffold, showing sparse cell attachment primarily along fiber grooves and at junctions. (B2-B3, C2-C3) The 60° and 45° scaffolds, showing increased cell numbers preferentially anchored to the higher density of fiber intersections and “edges.” (D2-D3) The 30° scaffold, exhibiting the highest cell density, but still failing to form a confluent monolayer, with cells instead clustering on the complex topography.

The SEM micrographs provide a stark and definitive confirmation of our hypothesis. While cells were able to sparsely attach to the PCL, their behavior was indicative of a highly unfavorable surface. Instead of forming a confluent, well-spread monolayer, the cells exhibited reluctant and highly selective attachment.

90° Scaffold (Fig. 9A1-A3): On the simple orthogonal grid, cell attachment was extremely limited. The few cells present were typically found stretched taut along the micro-grooves of individual fibers or clustered precariously at the fiber junctions (Fig. 9A2, A3). The vast, flat surface of the underlying substrate between the fibers remained almost entirely cell-free, highlighting the cells’ aversion to the hydrophobic material.

60° and 45° Scaffolds (Fig. 9B1-C3): These designs, featuring a higher density of fiber intersections, showed a noticeable increase in the number of attached cells compared to the 90° design. However, the attachment pattern remained suboptimal. Cells demonstrated a clear preference for topographical features, anchoring primarily to the “edges” provided by the fiber crossings and often spanning across open pores as suspended bridges (Fig. 9B3, C2, C3). This behavior suggests the cells were seeking points of high surface energy to secure attachment, rather than uniformly colonizing the surface.

30° Scaffold (Fig. 9D1-D3): This design, with the highest density of edges and intersections, correspondingly showed the highest density of attached cells among the unmodified groups. Cells were found extensively wrapping around and anchoring to the complex fiber network (Fig. 9D2, D3). Crucially, despite the higher number of cells, they still failed to form a confluent layer and instead existed as isolated clusters tethered to the PCL framework.

### 3.9. Live/Dead, Metabolic Activity, and Immunofluorescence Analysis of hTERT-HM Cell Interaction with MEW 3D-Printed constructs

Given the demonstrated superior wettability and tunable physicochemical properties of the composite film in comparison to unmodified PCL, a comprehensive series of in vitro studies was subsequently performed to comprehensively analyze and determine its overall biological performance.

Before evaluating the final composite scaffold, the cytocompatibility of its individual components and benchmark materials was assessed using a Live/Dead assay. This assay stains living cells green with Calcein AM and dead cells red with Propidium Iodide (PI). The results obtained from these control substrates and cell structures provide a crucial baseline for interpreting the performance of the final engineered construct (Fig. 10). hTERT-HM cells cultured on standard tissue culture plastic (Fig. 10A1-A3) served as the positive control. The images show a dense, confluent monolayer of cells. The vast majority of cells stain intensely green (Calcein), indicating high viability and robust metabolic activity. Only a very small, negligible number of red-staining (PI) dead cells are present. This confirms the intrinsic health and viability of the hTERT-HM cell line under standard culture conditions.

**Figure 10:**
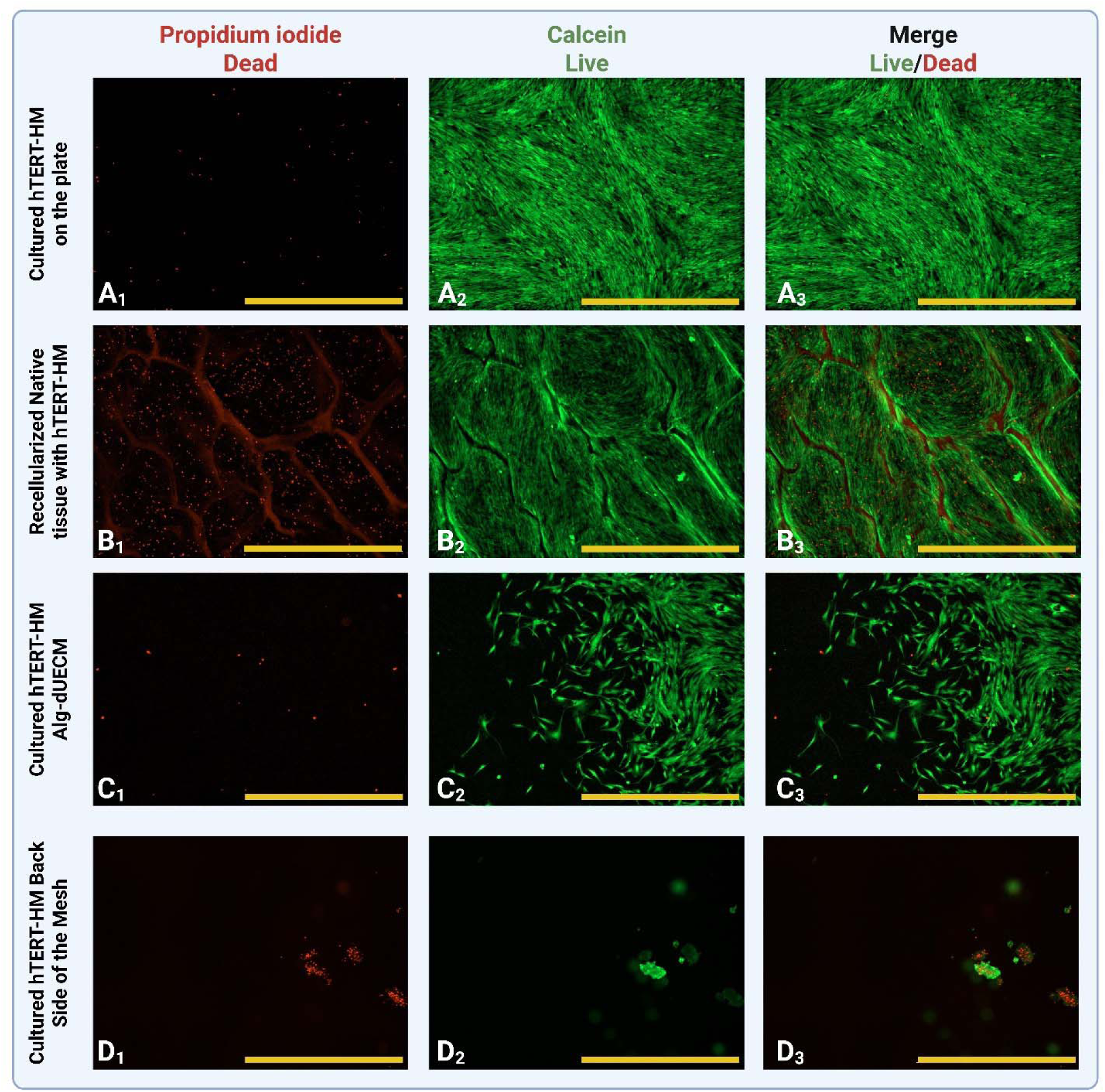
Live/Dead cytotoxicity assay of hTERT-HM cells on various control substrates after 7 days of culture. Live cells are stained green (Calcein AM), and the nuclei of dead cells are stained red (Propidium Iodide). (A1-A3) Positive control showing a dense, highly viable monolayer of cells on standard tissue culture plastic. (B1-B3) “Gold standard” control of cells seeded on recellularized native uterine tissue, showing high viability but with a notable baseline level of dead cells. (C1-C3) Hydrogel control showing excellent cell viability and spreading on a casted Alg-dUECM gel, confirming the cytocompatibility of the hydrogel component. (D1-D3) Cell death was significantly higher on the unmodified, hydrophobic, nonwoven PCL fibers arranged on the non-adhesive, back side of the membrane; these PCL fibers were nonwoven and distributed across the mesh’s reverse side. Scale bars = 1000 µm.

Native uterine tissue that was decellularized and then recellularized with hTERT-HM cells (Fig. 10B1-B3) served as the “gold standard” biological benchmark. The results show excellent cell viability, with a dense population of green-staining live cells repopulating the native ECM structure. However, a noticeable population of red-staining dead cells is also present, scattered across the tissue surface and along the native matrix ridges. This suggests that while native ECM is highly cytocompatible, the process of recellularization or the complex topography may induce a baseline level of cell stress or death.

To isolate the effect of the hydrogel component, cells were cultured directly on a casted Alg-dUECM gel without the PCL framework (Fig. 10C1-C3). The images reveal excellent cytocompatibility. The surface supports the attachment and spreading of hTERT-HM cells, which exhibit a healthy, elongated morphology and stain predominantly green. The number of dead cells is remarkably low, comparable to the positive control on the tissue culture plate. This critically demonstrates that the Alg-dUECM hydrogel itself is a highly permissive and cytocompatible substrate for hTERT-HM cells.

A key design feature of a barrier film is to prevent unwanted cell adhesion on its outer surface. The test of the unmodified nonwoven PCL fibers on the “back side” of the mesh provides crucial insight into this function (Fig. 10D1-D3). The images reveal a surface that is profoundly non-adhesive to hTERT-HM cells. Instead of forming a layer, the vast majority of cells appear to have failed to attach and were washed away during sample preparation. The only cells remaining are those that have managed to find micro-topographical niches. This leads to two critical observations: (A) The few surviving cell clusters are found precisely where the micro-scale grooves and curvatures of the PCL fibers can physically entrap cells or concentrate serum proteins from the media, creating isolated islands of permissive substrate. This confirms the hypothesis that the cells are not adhering to the bulk PCL material but rather to these specific topographical and protein-coated niches. (B) In these few aggregates where cells did manage to loosely attach, a high proportion of them are stained red (dead). This cell death is not indicative of material toxicity, but rather of anoikis—a form of programmed cell death that occurs when anchor-dependent cells fail to establish secure, widespread adhesion to a substrate [33].

Having established the essential role of hydrogel in promoting cell adhesion, the long-term biological performance of the final composite scaffolds was assessed. The viability and proliferative behavior of hTERT-HM cells on the different architectural designs (90°, 60°, 45°, and 30°) were analyzed at Day 1, Day 5, and Day 7 using a Live/Dead assay (Fig. 11).

**Figure 11:**
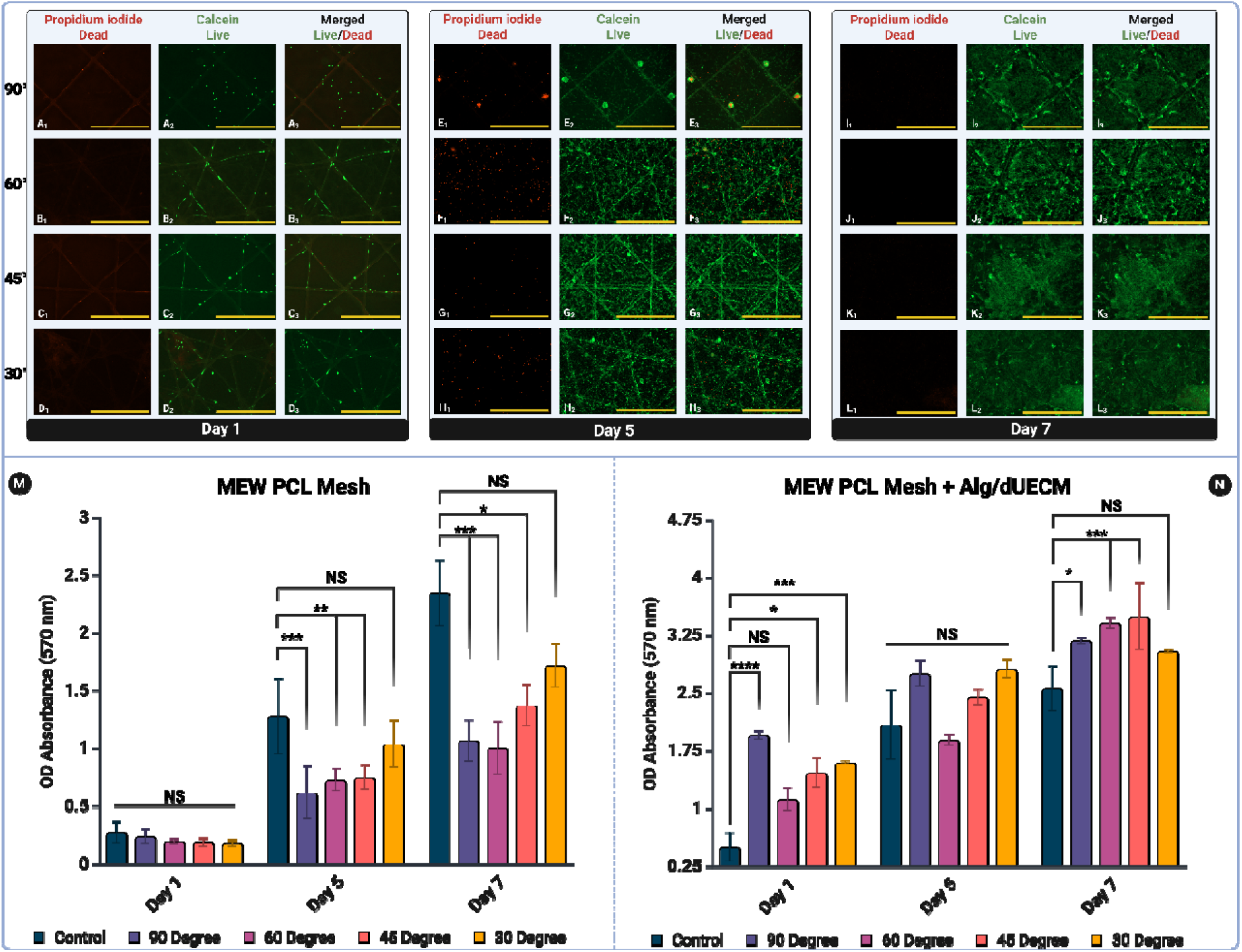
Comprehensive analysis of cell viability and proliferation, demonstrating the critical role of Alg-dUECM hydrogel in transforming the PCL mesh into a highly cytocompatible and bioactive scaffold. Top Panel (A-L): Qualitative time-course Live/Dead assay of hTERT-HM cells cultured on the PCL/Alg-dUECM composite scaffolds. Live cells are stained gree (Calcein), and dead cells are stained red (Propidium Iodide). (A-D) At Day 1, all scaffold architectures support high initial cell viability and attachment. (E-H) At Day 5, robust cell proliferation is evident across all designs. (I-L) By Day 7, a dense and highly viable confluent monolayer has formed on all composite scaffolds (Scale bars = 1000 µm). Bottom Panel (M, N): Quantitative MTT assay measuring the metabolic activity (OD Absorbance at 570 nm) of hTERT-HM cells over 7 days. (M) MEW PCL Mesh: Results from cells cultured on the unmodified PCL-only mesh. The data show very low initial activity and subsequent slow, inefficient proliferation, confirming the non-permissive nature of the pure PCL surface. (N) MEW PCL Mesh + Alg/dUECM: Results from cells cultured on the final composite scaffold. The data demonstrate robust, sustained proliferation from Day 1 to Day 7 across all architectures, with all groups significantly outperforming both the unmodified PCL and the tissue culture plastic control by Day 7. Statistical significance is denoted as *p < 0.05, **p < 0.01, ***p < 0.001, ****p < 0.0001, and NS (not significant). All data are presented as mean ± SD.

At just 24 hours post-seeding, the profound effect of the Alg-dUECM hydrogel was immediately apparent. All four composite scaffold architectures supported excellent initial attachment and high cell viability (Fig. 11A-D). The images show fields of viable, green-staining cells distributed across the scaffold surfaces. Critically, there are very few red-staining dead cells, confirming that the composite surface is highly cytocompatible and non-toxic from the outset. The underlying scaffold topography still influences initial cell distribution. On the 90° design (Fig. 11A2), cells appear more rounded and individually settled. On the more complex 60°, 45°, and 30° designs, cells begin to align and elongate along the denser fiber networks, demonstrating early-stage cell-topography interaction.

Day 5 images provide clear evidence of robust cell proliferation across all scaffold architectures (Fig. 11E-H). The density of green, viable cells has increased dramatically since Day 1, indicating that the composite surface actively supports cell division. The number of red-staining dead cells remains negligible. A key development is the transition from individual cells to an interconnected network. The hTERT-HM cells have begun to spread, elongate, and form connections with neighboring cells, creating a web-like sheet that spans the pores of the scaffold. This is particularly evident in the 60° (Fig. 11F2), 45° (Fig. 11G2), and 30° (Fig. 11H2) designs, and signifies the initial formation of a tissue-like layer.

By Day 7, the hTERT-HM cells successfully reached the desired outcome on all scaffold designs: the formation of a dense, confluent, and highly viable monolayer (Figure I-L).

By Day 7, the hTERT-HM cells successfully reached the desired outcome on all scaffold designs; the formation of a dense, confluent, and highly viable monolayer (Fig. 11I-L). The micrographs are dominated by an intense green fluorescence from a contiguous sheet of healthy, metabolically active cells. The underlying scaffold architecture is almost entirely masked by this complete cellular coverage. The near-total absence of red dead cells across all designs at this lat time point provides definitive evidence for the long-term cytocompatibility and health-promoting nature of the PCL/Alg-dUECM composite material.

To quantitatively support the qualitative observations from the Live/Dead assay, the metabolic activity and proliferation of hTERT-HM cells were measured using an MTT assay at Day 1, 5, and 7. Crucially, the assay was performed on two sets of scaffolds: the unmodified PCL-only mesh to establish a baseline (Fig. 11M), and the final PCL/Alg-dUECM composite scaffold (Fig. 11N). This direct comparison was designed to quantify the functional benefit of the hydrogel impregnation.

The MTT results from the unmodified PCL meshes reveal a surface that is not conducive to long-term cell growth (Fig. 11M). At Day 1, the metabolic activity on all scaffold groups was extremely low, reflecting the poor initial cell attachment to the hydrophobic surface. The mean OD absorbance values were just 0.24 ± 0.07 for the 90° design, 0.20 ± 0.02 for 60°, 0.19 ± 0.04 for 45°, and 0.18 ± 0.03 for 30°, all significantly lower than the control. Over the next several days, a trend of very slow and inefficient proliferation was observed. By Day 5, the mean OD values had modestly increased to 0.62 ± 0.22 (90°), 0.73 ± 0.09 (60°), 0.75 ± 0.10 (45°), and 1.04 ± 0.20 (30°). This slow growth continued to Day 7, with final mean OD values reaching 1.07 ± 0.18 (90°), 1.01 ± 0.23 (60°), 1.38 ± 0.18 (45°), and 1.72 ± 0.19 (30°).

While this data indicates that a small number of initially attached cells are able to slowly divide, it confirms that the unmodified PCL mesh is an inadequate substrate that cannot support the robust, exponential growth required for tissue engineering applications.

The MTT data from the final composite scaffolds tells a completely different and compelling story of success (Fig. 11N). At Day 1, all composite scaffold groups already exhibited significantly higher metabolic activity than the tissue culture plastic control (OD = 0.51 ± 0.19). The mean OD values were 1.95 ± 0.05 for 90°, 1.12 ± 0.15 for 60°, 1.47 ± 0.19 for 45°, and 1.60 ± 0.02 for 30°. This confirms that from the very first day, the hydrogel surface provides a highly permissive environment. From Day 1 to Day 7, a clear and consistent trend of robust cell proliferation was observed across all composite scaffold designs. For every architecture, the OD absorbance value significantly increased at each successive time point. For instance, the metabolic activity of the 45° scaffold increased from a mean OD of 1.47 ± 0.19 on Day 1, to 2.45 ± 0.10 on Day 5, and finally to 3.50 ± 0.43 on Day 7. By Day 7, the metabolic activity on all composite scaffolds was not only vastly superior to the struggling cultures on the unmodified PCL, but was also significantly higher than the tissue culture plastic positive control (OD = 2.56 ± 0.29). The final mean OD values were 3.19 ± 0.04 for 90°, 3.42 ± 0.07 for 60°, 3.50 ± 0.43 for 45°, and 3.05 ± 0.02 for 30°. The 60° and 45° designs, in particular, demonstrated the highest levels of cell metabolic activity, suggesting these architectures may provide an optimal microenvironment to enhance proliferation. The comparative and detailed MTT results are given in Table 5.

**Table 5:** Metabolic activity assessed by MTT (OD ^−--^ nm) (mean ± SD, n = 4; one-way ANOVA + Tukey within each day).

| Day | Plate | $90^\circ$ | $60^\circ$ | $45^\circ$ | $30^\circ$ |
| --- | --- | --- | --- | --- | --- |
| **Unmodified PCL Mesh** |  |  |  |  |  |
| **Day 1** | $0.28 \pm 0.10$ | $0.24 \pm 0.07$ | $0.20 \pm 0.02$ | $0.19 \pm 0.04$ | $0.18 \pm 0.03$ |
| **Day 5** | $1.28 \pm 0.32$ | $0.62 \pm 0.22$ | $0.73 \pm 0.09$ | $0.75 \pm 0.10$ | **$1.04 \pm 0.20$** |
| **Day 7** | $2.35 \pm 0.29$ | $1.07 \pm 0.18$ | $1.01 \pm 0.23$ | $1.38 \pm 0.18$ | **$1.72 \pm 0.19$** |
| **Impregnated PCL Mesh with Alg-dUECM** |  |  |  |  |  |
| **Day 1** | $0.51 \pm 0.19$ | $1.95 \pm 0.05$ | $1.12 \pm 0.15$ | $1.47 \pm 0.19$ | $1.60 \pm 0.02$ |
| **Day 5** | $2.09 \pm 0.45$ | $2.76 \pm 0.16$ | $1.89 \pm 0.07$ | $2.45 \pm 0.10$ | $2.81 \pm 0.12$ |
| **Day 7** | $2.56 \pm 0.29$ | $3.19 \pm 0.04$ | $3.42 \pm 0.07$ | $3.50 \pm 0.43$ | $3.05 \pm 0.02$ |

To provide definitive high-resolution visual evidence of the successful fabrication of the composite and its ability to support a confluent cell layer, SEM was conducted at each key stage of the assembly process.

The initial stage involved imaging the acellular PCL scaffolds (Fig. 12, Top Row). High magnification images (Fig. 12A1-D1) confirm the high-fidelity fabrication of the four distinct MEW architectures (90°, 60°, 45°, and 30°) on top of the nonwoven fiber base layer. High magnification views (Fig. 12A2-D2) reveal the precise, ordered MEW fibers and the underlying, more chaotic nonwoven fiber network. The back side of the scaffold (Fig. 12E1-E2) shows a highly porous, non-woven fibrous mesh designed to be a non-adhesive barrier.

**Figure 12:**
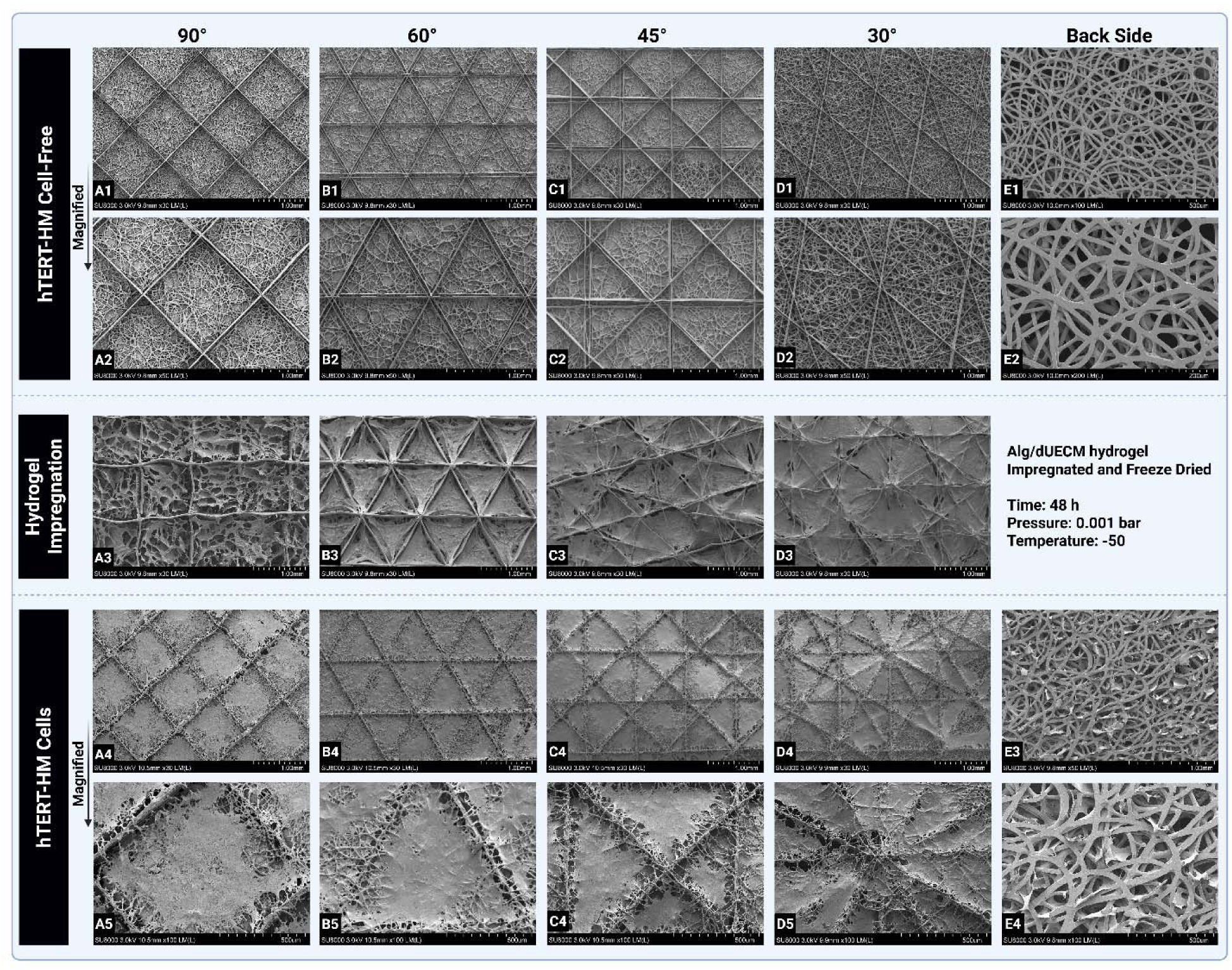
SEM analysis demonstrating the step-by-step fabrication of the composite barrier film and the successful formation of a confluent hTERT-HM cell monolayer. Top Row (Cell-Free): Micrographs of the acellular scaffold, showing the high-fidelity MEW architectures (90°, 60°, 45°, 30° in A1-D2) on top of a nonwoven fiber base layer, and the porous nonwoven mesh on the back side (E1-E2). Middle Row (Hydrogel Impregnation): Micrographs after impregnation with Alg-dUECM hydrogel and freeze-drying. A continuous hydrogel film is seen spanning the pores of the scaffold (A3-D3). The hydrogel has diffused through the entire construct and is mechanically entangled in the nonwoven fiber backing (E3-E4). Bottom Row (hTERT-HM Cells): Micrographs after 7 days of cell culture on the composite scaffolds. Low-magnification views (A4-D4) show the formation of a dense, confluent cell monolayer. High-magnification views (A5-D5) reveal healthy, well-spread cells completely covering the composite surface, creating a seamless biological barrier.

Following freeze-drying, the hydrogel-impregnated scaffolds were imaged (Fig. 12, Middle Row). The SEMs (Fig. 12A3-D3) clearly show that the Alg-dUECM hydrogel has successfully infiltrated the porous PCL framework. It forms a continuous, web-like film that spans the pores of the MEW grid and encapsulates the PCL fibers. This confirms the formation of an integrated composite material. Critically, imaging the back side of this composite (E3-E4) reveals that the hydrogel has diffused through the entire scaffold thickness to the nonwoven fiber layer. The hydrogel is seen to be mechanically entangled within the porous structure of the nonwoven mesh, demonstrating a strong physical interlock between the hydrogel and the PCL framework throughout the entire construct.

The final and most important stage was the analysis of the scaffolds after being seeded with hTERT-HM cells for 7 days (Fig. 12, Bottom Row). The results provide stunning visual confirmation of the success of the composite design. Low magnification images (Fig. 12A4-D4) show that the hTERT-HM cells have formed a dense and confluent monolayer that covers the entire surface of all scaffold architectures. The underlying grid pattern is still discernible, but the surface is now dominated by a contiguous sheet of biological tissue.

High magnification views (Fig. 12A5-D5) provide exquisite detail of the cell-material interface. The cells exhibit a healthy, flattened, and well-spread morphology, forming extensive cell-cell junctions and completely engulfing the underlying hydrogel and PCL fibers. They create a seamless biological barrier that bridges the pores of the scaffold. The surface topography is no longer that of a synthetic material but rather that of a living tissue layer.

To provide a final, functional assessment of cell behavior, the morphology and cytoskeletal organization of hTERT-HM cells were visualized using immunofluorescence staining for α-SMA (a key contractile protein; red/orange) and DAPI (to stain cell nuclei; blue). The analysis compares the composite scaffolds to control surfaces at both Day 1 and Day 7 of culture.

Cells on the standard tissue culture plate served as the positive control, showing well-spread individual cells with clearly defined nuclei and nascent actin filament networks (Fig. 13A1). At Day 1, hTERT-HM cells on all four composite scaffold architectures (90°, 60°, 45°, and 30°; Fig. 13B1-B4) demonstrated successful attachment and the initial formation of actin cytoskeletons. The cells began to adopt an elongated, spindle-like morphology, aligning with the local topography provided by the underlying hydrogel and PCL fibers. This early-stage organization is a key indicator that the cells are actively engaging with and responding to the scaffold’s surface.

**Figure 13:**
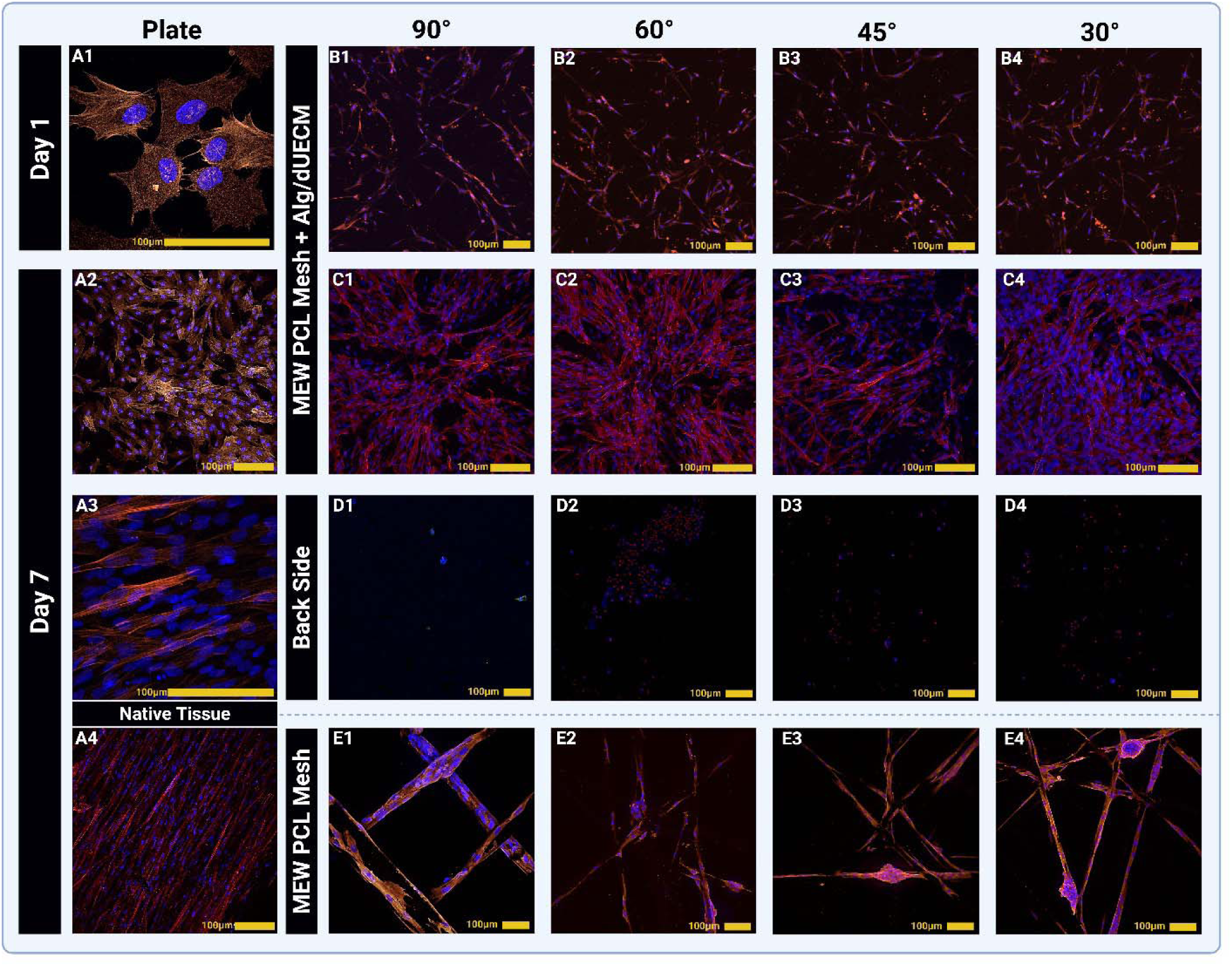
Immunofluorescence analysis of cytoskeletal organization, confirming the formation of a mature and contractile hTERT-HM cell monolayer on the final composite barrier film. Cell nuclei were stained with DAPI (blue), and the contractile cytoskeleton was stained for α-SMA (red/orange). Day 1 (Top Rows): (A1) Cells on the tissue culture plate control show initial, healthy spreading. (B1-B4) Cells on all composite scaffold architectures (90°, 60°, 45°, and 30°) demonstrate successful attachment and the formation of nascent α-SMA filaments, indicating active engagement with the substrate. Day 7 (Bottom Rows): (A2-A3) At Day 7, cells on the plate control (A2) have formed a dense but largely unorganized monolayer, while cells on the native tissue control (A3) show a highly aligned, physiological morphology. (C1-C4) Critically, cells on all four composite scaffold architectures have proliferated to form a dense, confluent monolayer. They exhibit robust expression of α-SMA organized into prominent, interconnected stress fibers, indicating the development of a mature, contractile tissue layer. (D1-D4) In stark contrast, the Back Side of the composite mesh is shown to be functionally non-adhesive, with virtually no cells present, confirming its barrier function. (E1-E4) As a final negative control, cells on the unmodified MEW PCL Mesh without hydrogel show sparse attachment, abnormal morphology, and a clear failure to form a monolayer, with cells clinging only to topographical features. Scale bars = 100 µm.

After 7 days of culture, a profound maturation of the cell layer was observed, highlighting the success of the composite material. The cells on the culture plate formed a dense, but largely disorganized, “cobblestone” monolayer (Fig. 13A2). In stark contrast, cells recellularizing native uterine tissue adopted a highly aligned, elongated morphology, with robust α-SMA expression organized into parallel stress fibers (Fig. 13A4). This represents the “gold standard” physiological organization for contractile smooth muscle tissue. By Day 7, the cells on all four composite scaffold designs (Fig. 13C1-C4) had proliferated to form a dense, confluent monolayer. Crucially, they exhibited robust expression of α-SMA organized into prominent, interconnected stress fibers. This indicates that the cells are not only viable but have differentiated into a mature, contractile phenotype. The organization of these fibers appears influenced by the underlying scaffold architecture, with cells on the more complex 45° and 30° designs (Fig. 13C3, C4) showing a more intricate, multi-directional network of actin fibers compared to the more orthogonal arrangement on the 90° design (Fig. 13C1).

The importance of the composite design is underscored by the negative controls at Day 7: As hypothesized, the non-adhesive PCL backing supported virtually no cell attachment. Only a few isolated, rounded cells with no discernible actin organization are visible, confirming its function as a cell-repellent barrier (Fig. 13D1-D4. Consistent with all previous findings, cells seeded on the pure PCL mesh failed to thrive. The few that remained were sparsely attached, highly elongated, or clumped and demonstrated a clear preference for topographical niches like fiber junctions and grooves. They failed to form a monolayer, and their actin expression was disorganized, indicating a stressed and non-functional state (Fig. 13E1-E4).

## 4. Discussion

Uterine tissue engineering presents unique biomechanical and biological challenges due to the multilayered structure of the uterus, each with distinct ECM organization and uterine smooth muscle function. In this study, we designed a bilayer scaffold that mimics both the outer serosal and the outer myometrial layers using a rationally engineered architecture that integrates directional fiber alignment and ECM bioactivity with surface hydrophilicity to support both regeneration and anti-adhesion outcomes.

As shown in Fig. 1A, the jet lands at a Jet contact point that lags behind the overhead nozzle, creating a characteristic jet angle (θj). The stability of this entire process is dictated by the relationship between the polymer jet’s extrusion velocity (V*_jet_* or V*_head_*) and the critical translation speed (V*_CTS_*), as conceptualized in the lower diagrams. Our optimization experiments identified V*_head_* = 70 mm s^−1^ as an ideal speed, corresponding to a regime where V*_CTS_* > V*_jet_*. This condition ensures that a constant drawing force is applied to the jet, pulling it straight and overcoming the inherent buckling instabilities (V*_jet_* > V*_CTS_*) that lead to fiber coiling at slower speeds.

By leveraging the optimized and stable printing process (V*_CTS_*> V*_jet_*), the capability to fabricate complex, multi-angled scaffold geometries was demonstrated (Fig. 1B). Using optimal parameters, scaffolds with programmed strand lay-down angles of 90°, 60°, 45°, and 30° were fabricated.

The optimized conditions yielded uniform fibers with minimal beading or jet instabilities, enabling precise layer-by-layer deposition. This observation aligns with prior MEW studies showing that collector translation speed is a dominant factor governing fiber diameter and pattern fidelity [34–36]. In one study, they reported that increasing collector speed significantly reduces fiber thickness, whereas overly slow speeds cause fiber stacking and inconsistency [37]. In our experiments, we also discovered a “Goldilocks” zone of speed and pressure parameters which facilitated the uninterrupted production of approximately 10 µm fibers without any structural failures; this finding is consistent with the optimal balance of flow rate (syringe pressure), voltage, Z-offset, temperature and speed for stable jet formation, as identified by other researchers’ response surface analysis [38]. In another study, they designed scaffolds with regular pore geometry (∼200–300 µm spacing), which is noteworthy given that pore size profoundly affects cell response [39]. They demonstrated that MEW scaffolds with ∼200 µm pores maximized the viability and matrix deposition of mesenchymal cells compared to those with smaller or larger pores. Thus, our printing optimization not only ensured structural precision but also created an architecture conducive to designing biomaterials that can mimic the complex anisotropic structure of native uterine tissue.

A key novelty of our work is the impregnation of the PCL scaffold with a composite hydrogel of Alg-dUECM. This biohybrid approach combines the mechanical strength and custom geometry of a synthetic scaffold with the biochemical complexity of a tissue-specific matrix, thereby addressing the limitations of each when used alone. Using decellularized uterine tissue provides important biological signaling cues [40]; however, this material is mechanically deficient and difficult to handle, a contrast to pure PCL meshes, which are strong but lack the necessary biological activity [41]. By combining them, we created a scaffold that is mechanically robust yet biologically instructive.

Physicochemical characterization confirmed successful integration of alginate and dUECM within the scaffold. FTIR spectra of the hybrid showed characteristic PCL ester bands (∼1720 cm^−1^) alongside new peaks around 1600–1650 cm^−1^ attributable to alginate’s carboxylate groups and the amide bonds of collagen/ECM proteins, which were absent in neat PCL. This is comparable to prior studies of PCL-natural polymer composites, where the inclusion of gelatin or ECM introduced amide I/II signals in the FTIR [42, 43]. Likewise, TGA revealed a multi-stage degradation profile: an initial drop around ∼260 °C (likely due to alginate and protein matrix decomposition) followed by the typical PCL thermal degradation above 350 °C. Such a profile mirrors other PCL/biopolymer systems, confirming that the dECM and alginate were effectively embedded in the scaffold. Importantly, the addition of alginate did not compromise PCL’s thermal stability; instead it was crucial for maintaining hydrogel presence. Recent studies have highlighted the crucial role of alginate addition in preserving the structural integrity of pure decellularized extracellular matrix (dECM) hydrogels throughout extended periods of cell culture [44]. Consistently, in our composite, alginate served as a crosslinkable backbone that gelled the soluble uterine ECM and prevented it from simply diffusing out of the scaffold. The Ca^2+-^crosslinked Alg-dUECM uniformly filled the PCL pores and adhered to fiber surfaces (as seen in SEM images in Fig. 12A3-D3), forming a stable bioactive coating throughout the construct.

Incorporation of the Alg-dUECM hydrogel dramatically improved the scaffold’s hydrophilicity and water absorption capacity. Pristine PCL is hydrophobic (water contact angle ∼100°) and resists wetting [45], which can limit protein adsorption and cell attachment. Our PCL scaffolds indeed showed high initial contact angles, but after Alg-dUECM impregnation the static contact angle dropped significantly (to ∼40–50°, a shift to hydrophilic behavior). This trend is fully consistent with literature on PCL/alginate composites. In a study, it was observed that the incorporation of alginate, at concentrations ranging from 10 to 40%, into PCL fibers resulted in a substantial improvement in their wettability and water absorption capabilities; specifically, these modified fibers exhibited water absorption levels up to twelve times greater than those observed in pure PCL fibers [46, 47]. Similarly, dynamic contact angle measurements by another researcher showed PCL/alginate nanofibers start around 92° but rapidly decrease to ∼45° as water permeates, whereas neat PCL stabilizes near 70° [48]. In our findings, the Alg-dUECM gel likely absorbed water and swelled within the pores, yielding a steady-state contact angle far below 70°, indicative of a highly wettable surface. The hydrophilicity augmentation is beneficial because moderate wettability promotes the adsorption of serum proteins and cell-adhesive molecules onto the scaffold, in turn facilitating cell attachment [49, 50]. Indeed, alginate and ECM components present hydrophilic hydroxyl, carboxyl, and amine groups that can bind integrin ligands from the dECM or recruit additional bioactive proteins from the culture medium.

In correlation with water retention, the Alg-dUECM scaffolds demonstrated swelling and degradation across all experimental groups. The early weight loss likely reflects partial alginate dissolution or release of ECM fragments not firmly crosslinked. While PCL by itself degrades very slowly (on the order of 3-4 years *in vivo*) [51], blending it with more labile components is known to accelerate scaffold resorption. For instance, blending PCL with faster-degrading PLA increased the overall degradation rate and resorbability of MEW scaffolds [45]. In our design, the alginate hydrogel phase can gradually disintegrate (especially in physiological fluids where ion exchange or alginate lyase may occur), and the dUECM proteins are susceptible to enzymatic breakdown. This creates porosity and channels that should allow subsequent cell-mediated degradation of PCL as healing progresses. The composite’s degradation behavior is thus favorable for tissue engineering: it provides an initial stable support, but becomes more porous and eventually resorbs over time, ideally matching the pace of new tissue formation. We acknowledge that PCL’s persistence is a potential limitation; however, by integrating degradable biological matrix we aimed to shorten the functional lifespan of the scaffold while still leveraging PCL’s mechanical support early on. Other researchers have pursued similar strategies of combining PCL with bioactive, faster-degrading phases to overcome PCL’s inertness and slow resorption. Our results reinforce those findings, showing that the Alg-dUECM hybrid scaffold achieves a balanced degradation profile suitable for uterine repair (where a scaffold may need to function for a few months to a year, but not remain indefinitely).

The improved swelling capacity of the composite scaffold also has implications for nutrient diffusion and cell survival. The Alg-dUECM hydrogel can hold a large amount of water within the scaffold, which should facilitate transport of oxygen and nutrients to cells even in the scaffold interior. In contrast, neat PCL is non-absorbent, and cells in its center can experience diffusion limitations. The swollen hydrogel matrix in our scaffolds likely helped overcome this, maintaining a hydrated environment that mimics native tissue.

Mechanical testing demonstrated that the hybrid PCL/Alg-dUECM scaffolds transitioned from a stiff, dry state to a more compliant and ductile form upon hydration, mimicking the mechanical behavior of native uterine tissue. This is attributed to water acting as a plasticizer and the deformable nature of the Alg–dUECM hydrogel, resulting the ductility of the MEW PCL/Alg-dUECM scaffolds. This mechanical softening in the hydrated state is advantageous for uterine applications, where the tissue must undergo cyclic expansion and contractile motion. The hybrid scaffold’s ability to deform harmoniously with the host tissue may minimize stress shielding and promote better integration. Although UTS in the wet state decreased, the dry scaffold remained robust enough for surgical handling. Importantly, by combining soft and fibrous phases, the composite structure approximated the mechanical compliance of myometrial tissue without relying on complex geometric modifications, representing a significant improvement over conventional rigid PCL meshes.

The biological performance of the Alg-dUECM scaffolds was significantly superior to that of plain PCL scaffolds, as evidenced by cell viability, proliferation, and phenotypic markers. Human uterine smooth muscle cells (hTERT-HM) readily adhered to the composite scaffolds and spread within the Alg-dUECM matrix, whereas on pure PCL fibers they showed sparse attachment and more rounded morphology (consistent with PCL’s bioinert surface). Live/Dead staining and MTT assays confirmed that the composite provided a cytocompatible niche: cell viability remained high over 7 days with metabolic activity substantially greater on Alg-dUECM scaffolds. In fact, by 7 days the cell number on composites had roughly doubled, whereas on PCL-only scaffolds it stagnated. This trend mirrors findings from other studies that combined PCL with natural biomolecules. For example, electrospun PCL nanofibers modified with a galactose ligand showed improved hydrophilicity and better support for cell attachment. This modification led to significantly higher viability and proliferation of uterine fibroblast cells compared to unmodified PCL fibers [52]. Our results are in strong agreement – the presence of the uterine ECM and alginate not only attracts more cells initially (higher seeding efficiency) but also supports their sustained growth. This can be attributed to the bioactive cues provided by the dUECM (such as collagen, laminin, growth factors from uterine tissue) which interact with cell receptors to promote attachment and proliferation. Additionally, the improved wettability allowed serum proteins (e.g., fibronectin) to adsorb, creating a conditioning layer for cells.

On the other hand, uterine tissue engineering must reproduce the uterus’s multilayered architecture while preventing post-operative adhesions. We therefore created a bilayer scaffold that merges (i) a random, hydrophilic PCL mesh that emulates the low-friction serosa and limits the adhesion of adjacent tissues, with (ii) an aligned (30°) MEW-PCL lattice impregnated with Alg-dUECM to recreate the contractile outer myometrium. dUECM supplies native collagens, fibronectin, GAGs, and bound growth factors, providing lineage-specific cues absent from inert barriers. This bioactivity was confirmed by abundant α-SMA staining in hTERT-HM cells cultured on Alg-dUECM constructs, whereas plain PCL showed minimal α-SMA. Hence, this composite scaffold not only blocks early adhesion formation but also preserves or induces the contractile smooth muscle cell phenotype, which is essential for functional myometrial regeneration.

In a study, researchers cultured menstrual blood-derived stem cells on a decellularized uterine scaffold and found that the cells differentiated into uterine lineage cells, expressing markers like cytokeratin (epithelial) and α-SMA within 10 days [53]. The decellularized uterus matrix acted as an instructive niche for lineage-specific differentiation [53]. Similarly, in our hybrid scaffolds, the dUECM component likely contains myometrial ECM proteins and bound factors that signaled the hTERT-HM to maintain their contractile phenotype. This is a crucial point of progress: rebuilding uterine tissue requires not just any cell growth but the correct cell phenotype (contractile smooth muscle in the myometrium) to restore function. Our work demonstrates that the composite scaffold can support this phenotypic stability, in line with other tissue-engineered uterus approaches using dUECM.

Thus, our biofabrication approach offers a promising solution for both uterine tissue engineering and post-operative anti-adhesion barrier applications. Traditional repair methods—such as decellularized uterine grafts or synthetic meshes—either lack mechanical strength or bioactivity, and often fail to prevent adhesion formation. Decellularized tissues integrate well but are limited by donor availability and structural control, while synthetic meshes may provoke chronic inflammation or fail to remodel with host tissue. Our melt-electrowritten PCL scaffold addresses these limitations by offering high-resolution, patient-specific designs with mechanical robustness. By incorporating uterine-derived ECM into the scaffold, we biologically functionalize the mesh, introducing regenerative and immunomodulatory signals known to reduce fibrosis and adhesion formation. This dual-function scaffold supports both structural repair and anti-adhesion functionality.

Unlike conventional electrospun mats, which are often too dense for sufficient cell infiltration and tend to act as passive barriers, our MEW scaffolds feature controlled pore sizes that permit deep cellular integration. The addition of a hydrogel layer enhances tissue compatibility and further acts as a physical barrier against postoperative adhesions by mimicking the hydrated, lubricative environment of the native serosa. By focusing on the myometrial layer, we demonstrate the platform’s potential, and future adaptations may enable a multilayered uterine wall reconstruction by adding endometrial-mimicking layers. The scaffold shows promise, as demonstrated by the in vitro results which revealed strong cell proliferation and α-SMA expression; however, additional research is necessary to confirm this potential. Our method integrates the structural precision of MEW with the bioactivity of dUECM, offering a new direction for functional uterine tissue engineering. Together, these features position our hybrid scaffold as a next-generation bioactive barrier for uterine repair with broad potential in gynecologic tissue engineering.

## 5. Conclusions and Future Directions

Despite recent advances in uterine regenerative therapies and anti-adhesion barrier designs, the clinical management of post-surgical adhesions remains challenging due to the lack of biomaterials that can simultaneously support tissue regeneration, restore biomechanical integrity, and prevent abnormal cell infiltration. Conventional synthetic barriers often fail to support myometrial or peritoneal regeneration due to their poor cytocompatibility, hydrophobic nature, and lack of uterine-specific signaling. Moreover, the unique microstructure of uterine tissue—with anisotropically aligned smooth muscle cells in the myometrium and randomly organized ECM in the serosal layer—necessitates a design that biomimics both architectural and biochemical features of the native uterus.

This study aimed to engineer a multilayered, biomimetic scaffold using near-field melt electrowriting of polycaprolactone integrated with a bioactive hydrogel composed of alginate and decellularized uterine extracellular matrix. The objective was to create a dual-function platform capable of acting as both an anti-adhesion barrier and a regenerative matrix to promote uterine scar healing. Our results showed that the composite scaffold not only overcame PCL’s intrinsic hydrophobicity but also exhibited superior cytocompatibility, tunable mechanical and degradation properties, and supported hTERT-HM cell adhesion, proliferation, and contractile phenotype expression. Techniques such as FTIR, TGA, SEM, wettability assays, and in vitro biological evaluations confirmed scaffold integration and function. The mechanical performance of the hydrated composite exceeded that of native uterine tissue, and its architecture could be modulated to match physiologic needs. This work provided a powerful platform technology for the development of advanced, bio-functional implants for uterine repair and other soft tissue engineering applications.

Future directions for this research should include in vivo evaluations within uterine injury or adhesion models, alongside comprehensive studies on long-term degradation and remodeling processes, and the exploration of integrating therapeutic molecules or primary patient-derived cells for enhanced efficacy and personalized treatment approaches. This platform offers strong translational potential not only for uterine repair but also for broader applications in soft tissue engineering where bioactive mechanical support and spatially organized cellular responses are required.

## 7. CRediT Authors Statement

**AFAY:** Conceptualization, Data curation, Formal analysis, Investigation, Methodology, Project administration, Software, Validation, Visualization, Writing – original draft, Writing – review & editing. **KT:** Writing–review and editing, Validation, Methodology, Conceptualization. **ME:** Writing–review and editing, Software, Formal Analysis, Data curation. **XT:** Data curation, **DJM:** Writing–review and editing, Validation, Resources. **IB:** Writing–review and editing, Validation, Supervision, Resources, Funding acquisition. **XC:** Writing–review and editing, Validation, Supervision, Resources, Funding acquisition.

## 8. Declaration of generative AI and AI-assisted technologies in the writing process

The author acknowledges Biorender.com for providing access to their platform for creating graphical illustrations. During the preparation of this manuscript, the authors used ChatGPT (GPT4o), Grammarly, and ProWritingAid for editing, grammar check, and clarity. The authors have reviewed and edited the output and take full responsibility for the content of this publication.

## 9. Institutional Review Board Statement

Not applicable.

## 10. Informed Consent Statement

Not applicable.

## 12. Data availability statement

The raw data supporting the conclusions of this article are available from the first and corresponding author upon reasonable request.

